# Efagins – engineerable agents that evolved independently to target the enterococcal cell wall

**DOI:** 10.64898/2025.12.13.693970

**Authors:** Janira Prichula, Abigail L. Manson, Joshua T. Smith, Natia Karumidze, Gracie Richards, Suelen S. Mello, Artur Czajkowski, Mohammed Kaplan, Ashlee M. Earl, Michael S. Gilmore

**Affiliations:** Department of Ophthalmology, Massachusetts Eye and Ear Infirmary, Harvard Medical School, Boston, MA, USA; Department of Microbiology, Harvard Medical School, Boston, MA, USA; Infectious Disease and Microbiome Program, Broad Institute of MIT and Harvard, Cambridge, MA, USA; Eliava Institute of Bacteriophages, Tbilisi, Georgia; European University, Tbilisi, Georgia; University of Chicago, Chicago, IL, USA

**Keywords:** *Enterococcus faecalis*, generalist species, efagin, phage-related elements, antimicrobial agents, Epa, Rha-CWPS, VRE, *Enterococcus faecium*, hospital acquired infection

## Abstract

Enterococci are major causes of multidrug-resistant infections, and antimicrobials with fundamentally new mechanisms of action are urgently needed. We identify a new class of antibacterial agents, termed efagins, which are chromosomally encoded, phage-related nanomachines that recognize cell wall carbohydrate receptors and inhibit subsets of *E. faecalis, E. faecium* and other enterococci selectively—a key reason they evaded prior detection. Five natural variants with distinct targeting profiles were identified **–** four related by sequence divergence, while one appears to have arisen through recombination of the targeting domain, likely from a phage donor. Operons encoding the corresponding carbohydrate receptors are highly variable, accounting for targeting specificity. The efagin targeting domain can be engineered to reprogram them toward alternative receptors, providing a pathway for filling critical coverage gaps. These findings advance efagins as new selective antibacterials with promise for addressing infection and spread of multidrug-resistant enterococcal infection.

## INTRODUCTION

Enterococci are projected to have evolved as gut microbes of land animals about the time arthropods first terrestrialized 400 – 500 MYA,^1^ but now rank among leading causes of multidrug-resistant (MDR) infections of humans worldwide.^2–4^ Of over 90 species present in diverse land animals^5^, two that are distantly related^1^ – *E. faecalis* and *E. faecium* – cause the vast majority of MDR human infection. Because of their ancient origins both naturally occur in non-human hosts, with *E. faecalis* being by far the greatest generalist of all enterococci.^5^ This raised the question as to what unique features of *E. faecalis* distinguish it from all other enterococcal species and allow it to successfully compete with more specialized enterococci and other species, and colonize diverse gut consortia as they are conveyed up and down the food chain.

Comparing >3,000 *E. faecalis* genomes from diverse ecologies, we observed that despite its ancient emergence and host diversification over eons, a 14.6 kb cluster of genes previously annotated as encoding a partial phage^6^ (which could not be ascribed phage function even in the presence of helper phages^7^) was highly conserved in all strains with reading frames well maintained. We interpret this as evidence that the product of this block of genes contributes fundamentally to the biology of *E. faecalis* in the highly diverse ecosystems in which it occurs.

Here, we show that this gene block encodes an independently evolved entity with antimicrobial activity that targets the rhamnose-rich enterococcal cell wall polysaccharide (Rha-CWPS) of enterococci and possibly other microbes. We term its product efagin (*E.faecalis* ph<u>ag</u>e-related <u>in</u>hibitor). Although sharing little sequence identity, efagins appear organizationally related to independently derived elements termed tailocins.^8,9^ Efagin activity extends beyond *E. faecalis* to include many *E. faecium* and other enterococci, likely contributing to the success of this species in colonizing diverse animal guts. The efagin targeting domain is amenable to reprogramming through domain swaps, highlighting a pathway for filling critical gaps in coverage and potentially targeting new organisms.

## RESULTS

### Efagins are encoded within all *E. faecalis* genomes from diverse ecosystems and are exclusive to the species

In a comparison of 3,068 genome sequences from diverse geographic and ecological sources, the efagin operon was found to be very highly conserved (Fig. 1, Table S1). All high-quality drafts (fewer than 75 contigs, n=2278) possess the entire element with reading frames intact; only five publicly available, low-quality draft genomes the lacked the full element did have partial hits. No *E. faecalis* genomes were found lacking efagin genes. The efagin gene cluster shows no evidence of mobility as its chromosomal neighborhood is highly conserved discounting any association with a mobile element (Extended Data Fig. 1). Efagin is therefore core to the *E. faecalis* species, and we interpret the high level of sequence conservation to reflect strong selection in diverse ecosystems.

**Fig 1.**
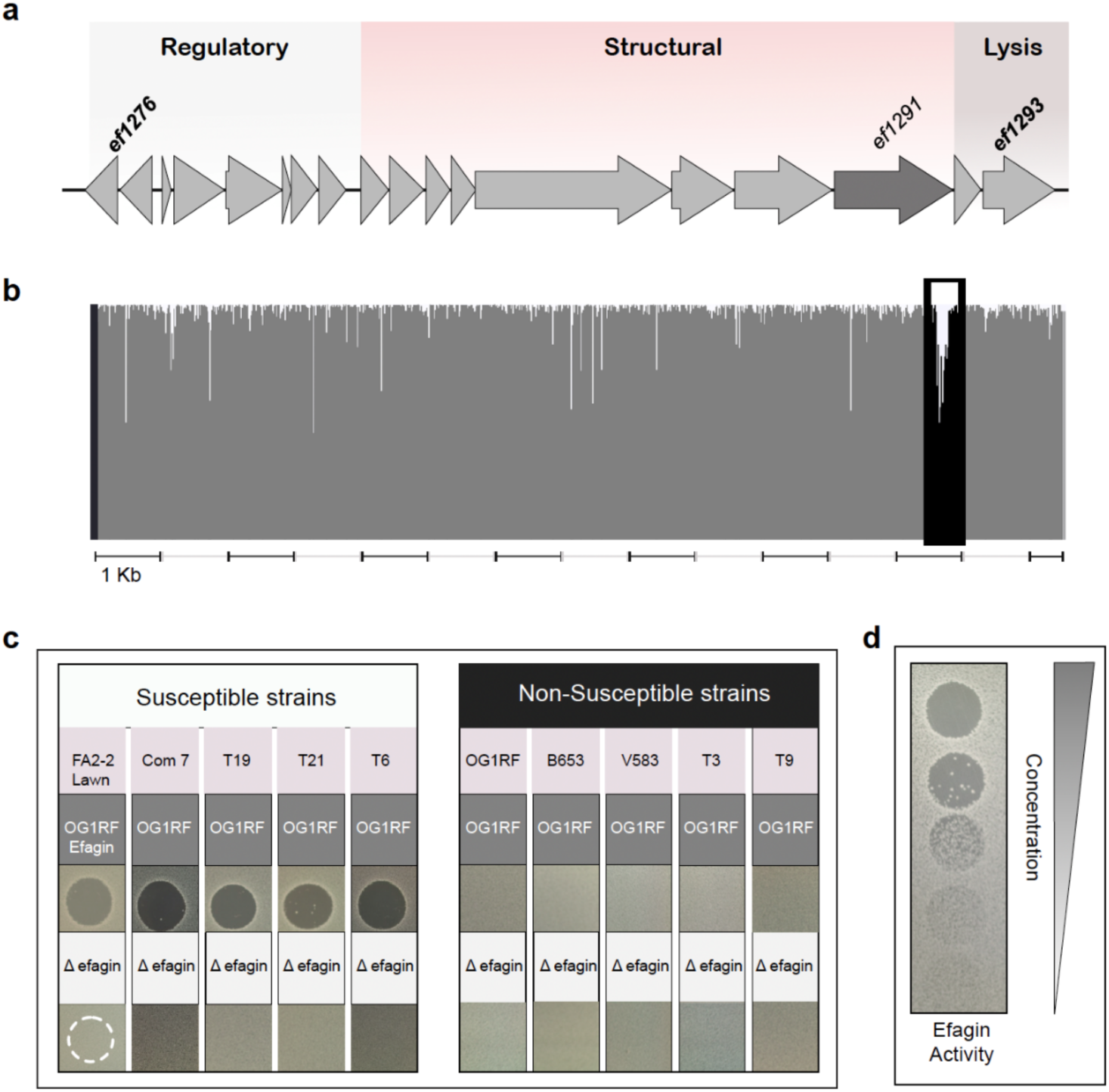
Efagin is core to *E. faecalis* and highly conserved except for a discrete variable region. **a,** Genetic organization of the 14.6kb efagin gene cluster, which in strain V583 consists of genes *ef1276* – *ef1293*. **b,** Alignment of corresponding sequences in 2,774 high quality *E. faecalis* genomes using MUSCLE^26^ shows the largest cluster of variability to occur in the C-terminal domain-encoding sequence of *ef1291* (highlighted). **c,** Efagin inhibits growth of select heterologous *E. faecalis* strains and not the producing organism (source of target cells [lavender box], source of efagin [OG1RF, top row; isogenic OG1RF(*Δefg*) bottom row). **d,** Efagin dilution does not result in discrete plaque formation but breakthrough growth instead.

To determine if efagin or anything analogous occurs in other enterococcal species, we searched a dataset of 15,854 *Enterococcaceae* genomes not including *E. faecalis*. No significant identity to efagin (>90% coverage; E-value <10) was found. A search of all draft prokaryotic genomes (approximately 1.2 million submissions) in NCBI (https://www.ncbi.nlm.nih.gov) similarly did not identify significant identities outside of *E. faecalis*. A broader search for: i) the presence of at least a partial phage, and ii) its occurrence in at least 50% of strains of a species (using the Genomad provirus detection tool^10^ with candidate prophage protein annotations confirmed using Phastest^11^) found that *E. faecalis* efagin was the only example fitting those criteria. Our search did identify an unrelated incomplete *Melissococcus plutonius* prophage present in 89.5% of the comparatively few genomes available for that species (17/19; Tables S2 and S3). However, its structural characteristics diverge from those of efagin as the *M. plutonius* element is predicted to encode a capsid and scaffold proteins suggestive of a defective phage or gene transfer agent^12^ (Table S3), in contrast to efagins, which are characterized by a complete absence of phage head-related genes (Fig. 1a; Extended Data Fig.2; Supplementary Fig. 1).

**Fig. 2.**
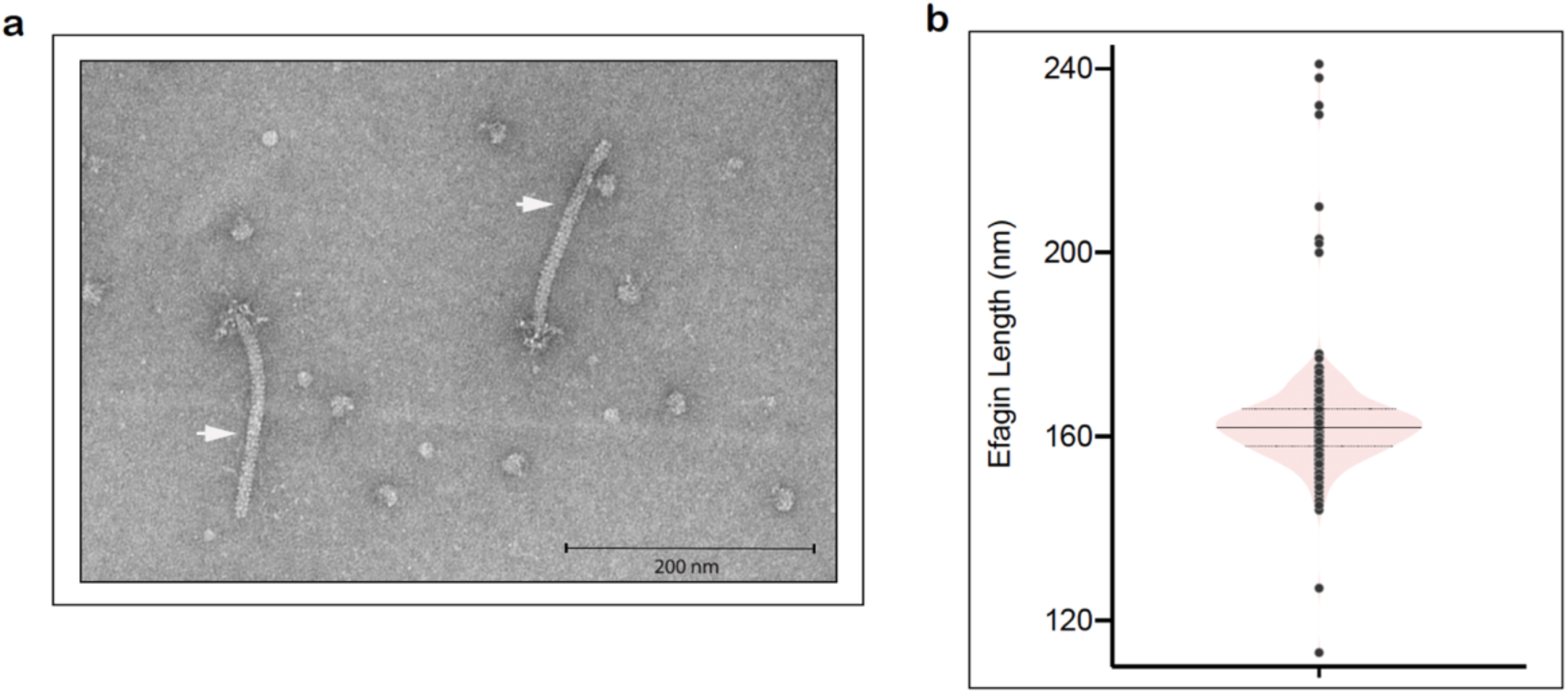
Efagin appears as a phage tail-like particle. **a,** TEM image of OG1RF efagin particles (white arrows) with phage tail-like structures including apparent baseplate. **b,** Size distribution of 300 efagin particles.

### Efagins are evolutionarily distant from Gram-positive and Gram-negative phage-related antimicrobial agents

Other antimicrobial phage-related systems have been identified.^13–24^ Though efagin shares little to no sequence identity with any, its genetic organization is broadly similar to the tailocin, monocin^20–22^ (Extended Data Fig. 2). At the amino acid sequence level, evidence of distant sequence identity is detectable only between three protein counterparts (putative regulatory element EF1276 exhibits 40% amino acid identity over 90% of its length with a monocin counterpart, whereas putative structural components EF1288 and EF1292 exhibit 21% identity over 59% of length, and 38% over 49% of the length respectively [Table S4]). For comparison, the ubiquity of monocin and other tailocins within genomes of the respective species that express them was examined and found to vary widely in contrast to the high degree of efagin conservation in *E. faecalis* (Tables S5-12; Extended Data Fig. 3). In current databases, *E. faecalis* phage vB_EfaS_IME197 (41.3 kb)^25^ exhibits the closest relationship – six efagin proteins share 24.6% to 70.7% amino acid sequence identity with its inferred gene products (Table S13), highlighting its independent evolution.

### Efagin is selectively antimicrobial

Previous work^7^ systematically explored seven putative prophage-related sequences moored within the prototype *E. faecalis* VRE V583 chromosome, including then-annotated partial phage-related element described here as efagin. All putative prophage-related sequences (alone or with added helper phage), <u>except</u> for efagin, produced viable particles. Separately, as a byproduct of work assessing vesicle formation by *E. faecalis*, others^27^ observed spurious copurifying particles which were attributed to genes of the efagin cluster, but the significance of those particles was not explored further.

Because efagin reading frames are well maintained and particles appeared to be expressed from them,^27^ we undertook a search for activity. We reproduced the observations of Afonina et al.^27^ by precipitating efagin exclusively from sterile culture filtrates of wild-type OG1RF and not from an isogenic strain generated here (OG1RF*Δefg*) in which the efagin gene cluster had been deleted, revealing phage tail-like particles.

To detect activity, paired sterile filtrates of OG1RF or OG1RF*Δefg* were assayed for potential antagonistic activity using a known diverse panel of *E. faecalis* lineages.^28^ Efagin-specific inhibition was observed for some *E. faecalis* but not others (Fig. 1c). Notably, efagin expressed by OG1RF did not kill the producer itself, but was active against select other *E. faecalis* strains. Furthermore, when serially diluted, efagin inhibitory activity was lost evenly (rather than forming fewer well-resolved individual plaques, as is typical of phages capable of self-replication and reinfection), with breakthrough growth at lower concentrations as seen with some antibiotics (Fig. 1d). Further, inhibition appeared confined to the spot of application on the lawn with little diffusion.

### Efagin structure

By transmission electron micrography (TEM), efagin particles measure from 113-234 nm, with a mean of 163 ± 11.8 nm (Fig. 2).

### *ef1291* Sequence variation is non-random

The observation of a concentrated patch of variability near the 3’ end of one gene (*ef1291*) suggested either a unique opportunity for sequence drift or selection for variation in the function encoded (Fig. 1b). We therefore examined it closely and found that all variants fell into one of five patterns, termed efagin types A-E (Fig. 3 and Table S14), supporting the latter prospect. Compared to (arbitrarily chosen) type A, variations in types B, C, and D clustered around specific amino acid positions. In contrast, the C-terminus of type E was unrelated (19.1% amino acid identity with type A) and appears to have arisen through allelic substitution – based on sequence similarity, possibly with a phage of the *Caudoviricetes* group (Table S15). We interpreted natural allelic replacement of this C-terminal domain to indicate that it likely encodes an independently functional domain. Additionally, a subvariant (which we named type X) was observed to occur within representatives of types B (2 subvariants), type C (49 subvariants), and type D (2 subvariants). Each of these subvariants possesses small sequence perturbations resulting in a premature stop codon that truncates EF1291 near the middle.

**Fig. 3.**
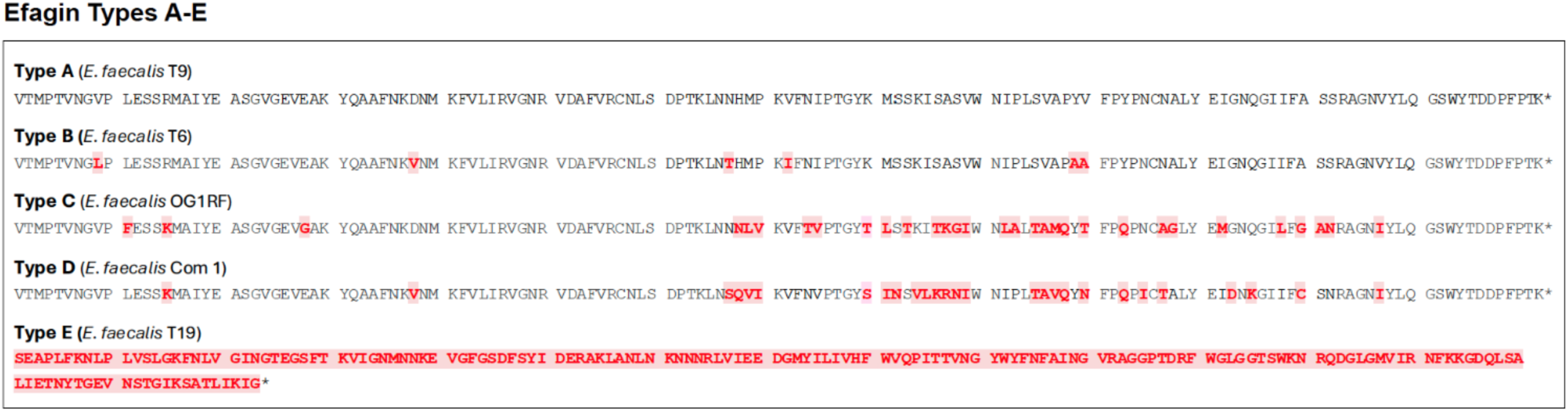
EF1291 C-terminal sequence variation patterns. Variations in encoded amino acids 448 to 619 fall into distinct classes defining efagin types A-E (differences from type A highlighted in red). Type E appears to have arisen from recombination.

### Structural similarity between EF1291 and a listeriophage carbohydrate-recognition domain

With limited inferred amino acid sequence identity to proteins of known function, we used Phyre2^29^ to detect possible conserved structural motifs (Table S16). Phyre2^29^ modeled 152 inferred C-terminal amino acids of efagin to the structure of the receptor-binding domain of Listeria phage PSA protein gp15 with 96.7% confidence. That domain of gp15 binds specific carbohydrate decorations on the variable *L. monocytogenes* wall teichoic acid (WTA).^30^ We therefore used AlphaFold^31^ to make a comparative efagin EF1291 model to map EF1291 sequence variations observed on its hypothetical structure (Extended Data Fig. 4). As listeriophage gp15 was solved in trimeric form,^30^ a quaternary structure common to other phage target binding proteins,^32–34^ EF1291 was modeled as a trimer as well. Variations defining efagin sequence types mapped to the positions of amino acids of gp15 including those known to contact specific carbohydrates of *Listeria* target cell WTAs^30^ (Fig. 4).

**Fig. 4.**
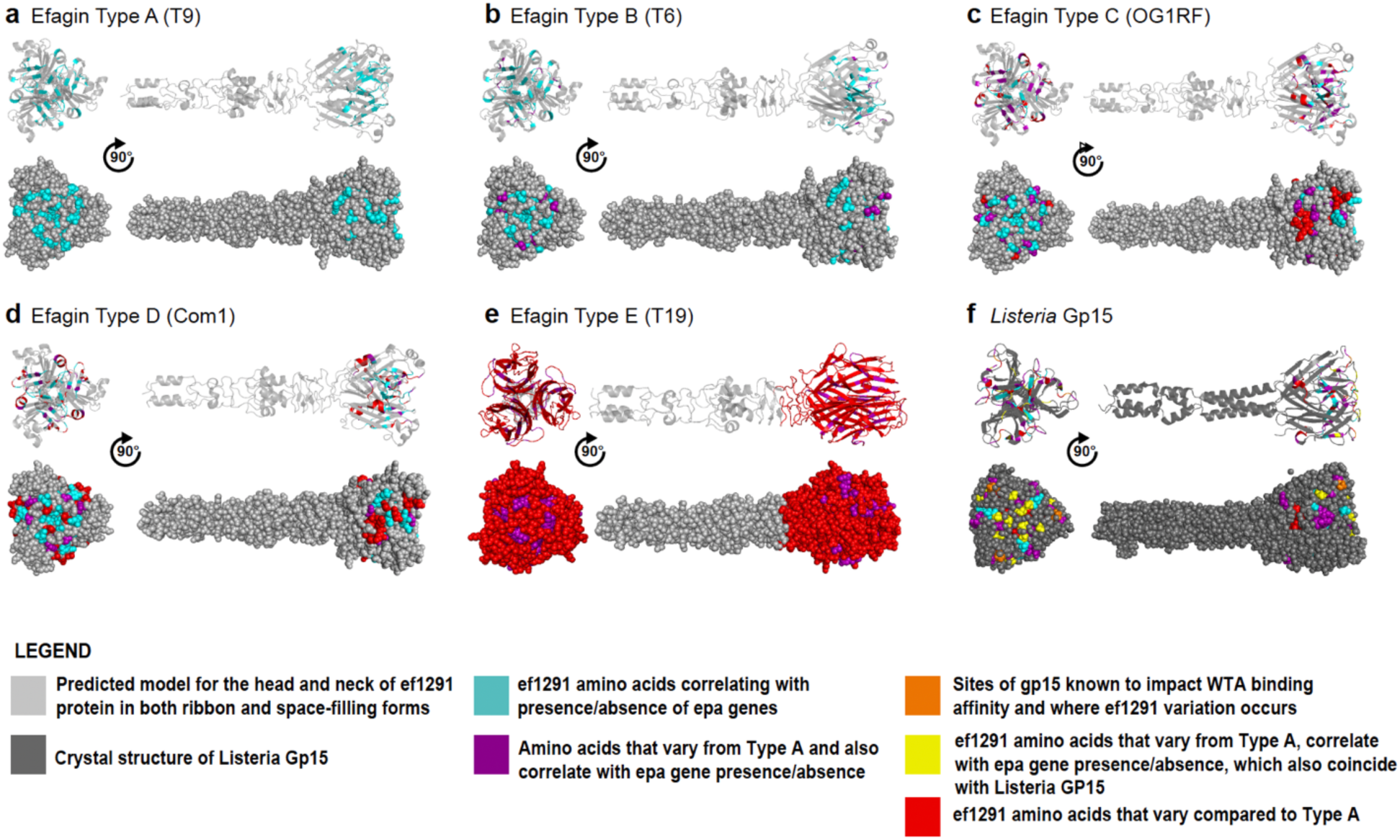
Structural similarities between EF1291 and listeriophage gp15 includes known carbohydrate-recognition domain. **a-e,** Base (left) and side (right) views of the predicted trimeric models of EF1291 in efagin types A-E using Alphafold 3.^31^ Amino acids that differ from efagin type A EF1291 are indicated in red. Amino acids that correlate with the presence/absence of *epa* genes (correlation coefficient < -0.6) are colored cyan. Amino acids that vary in both variation and correlation analysis are colored purple. **f,** Crystal structure of listeriophage PSA gp15^30^. Amino acids known to impact WTA binding in gp15 are colored yellow.^30^ Orange amino acids are those known to impact WTA binding in gp15^30^ and also vary in EF1291.

The rhamnose-rich cell wall polysaccharide (Rha-CWPS) referred to as enterococcal polysaccharide antigen (Epa)^35,36^, is the *E. faecalis* analog of the WTA targeted by listeriophage PSA protein gp15^30^ and is also known to vary between strains.^37,38^ Because the producing cell is not killed by its own efagin, we hypothesized that cells expressing various efagin types would possess different Epa cell wall carbohydrate structures to avoid self-targeting. We therefore tested covariation between the single nucleotide polymorphisms (SNPs) encoding the EF1291 efagin type variations, and the presence/absence of specific Rha-CWPS/*epa* operon genes indicative of alternate Epa structures, across the *E. faecalis* species diversity. Significant correlations were found between select *ef1291* SNP positions and the presence/absence of four genes in the *epa* cluster^36,38^: *ef2166* (-0.729, p=1.13e-91), *ef2167* (-0.729, p=1.13e-91), *ef2168* (-0.723, p=1.46e-89); and *e*f*2169* (-0.698, p=6.05e-81) (Extended Data Fig. 5 and Supplementary Fig. 2).

To experimentally verify Epa as the efagin target, we selected and sequenced efagin-resistant mutants of susceptible strains. Partially purified and concentrated type C efagin from OG1RF was spread onto lawns of susceptible *E. faecalis* strains FA2-2^39^ and X98.^40^ These targets were chosen because both are susceptible to type C efagin, but enigmatically were previously shown to be of different *epa* genotypes.^37^ Four FA2-2 mutants (FA2-2R1-R4) and two X98 mutants (X98R1 and -R2) were chosen for genome sequence comparison to parental controls. All type C efagin resistant mutants of both lineages possessed mutations in *epa* operon genes (Table S17 and Extended Data Fig. 6), providing direct experimental evidence that despite differences in *epa* genotype, wild type Epa is necessary for efagin activity.

Building on a prior analysis,^37^ we re-examined the diversity of *epa* operons in 1,972 high quality *E. faecalis* genomes. The *E. faecalis epa* cluster consists of 18 conserved core genes (*epa*A*–epa*R) followed by variable genes downstream (with *ef2164* to *ef2198* constituting the variable *epa* genes of prototype VRE *E. faecalis* strain V583)^36,37^ (Table S18). Therefore, the diversity of *epa* operons was clustered based on the extent of shared homologs in the variable gene set, which identified 20 *E. faecalis epa* genotypes (Fig. 5 and Table S14). All four genes whose presence or absence covaried with type-defining efagin variations (*ef2166*–*ef2169*) are widely shared but not universal in the variable gene set (Fig. 5).

**Fig. 5.**
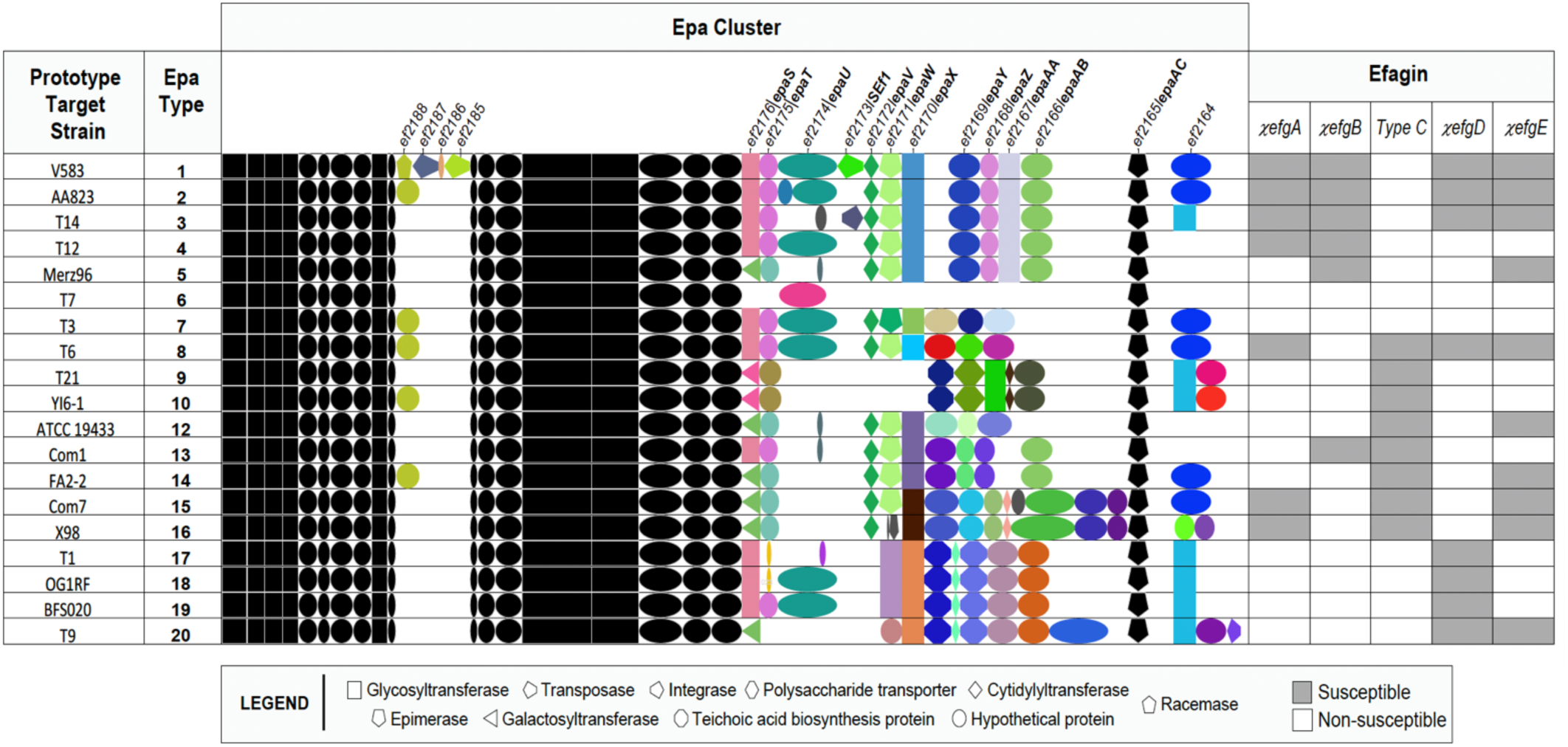
Epa genotypes and their susceptibility to targeting by efagin types. Representative operons of 19 of 20 identified *epa* genotypes occurring in strains available to us are symbolically represented (black [core] and colored [variable] shapes correspond to predicted function). Susceptibility to inhibition of each *epa* genotype strain to efagin types A-E is indicated (grey boxes), as measured using chimeras constructed on the OG1RF type C background to control for other potential antimicrobial activities potentially expressed by natural producers. Chimeras are labeled using the Greek letter “χ”(*chi*), followed by efagin (*efg*) type.

### The EF1291 C-terminal domain determines target cell specificity

To prove that the C-terminal domain of EF1291 determines target cell specificity, chimeras possessing C-terminal domain swaps from EF1291 sequence variation type A, B, D, and E donors, were constructed on the efagin type C background of OG1RF by allelic replacement.^41^ This allowed assessment of efagin activity and targeting free of other potential antagonistic activities or phages common in wild-type natural producers. Generation of an efagin type D EF1291 C-terminal chimera proved initially difficult. We reasoned that efagin type D may target OG1RF, making such a construct suicidal. Direct assessment of the supernatant of the type D (C-terminus donor strain Com1) supported this prospect (although other potentially Com1-expressed inhibitory activities were not controlled for). Nevertheless, using partially purified Com1 efagin to select for resistant OG1RF mutants, two independently derived colonies were selected for genome sequence analysis, OG1RF-R1 and OG1RF-R2. Consistent with expectation, each possessed a different mutation in the OG1RF *epa* operon variable region (Extended Data Fig. 6 and Table S17). Strain OG1RF-R2 was arbitrarily selected as the host for construction and expression of the type D chimera, which was obtained without further difficulty. Once obtained, the type D chimera was verified as targeting OG1RF, but not OG1RF-R1 and -R2 mutants. Each chimeric construction (termed χ*efg*A, B, D, and E, respectively) was obtained on the OG1RF type C efagin background and verified by sequencing. Construction of EF1291 chimeras did not detectably affect efagin assembly or structure (Supplementary Fig. 3)

Each efagin chimera was systematically examined for altered target cell specificity using representatives of 19 of the 20 *epa* genotypes identified (Fig. 5), as we were unable to obtain an *epa* genotype 11 strain for testing (Supplementary Fig. 4). Chimeric efagins χ*efg*A, χ*efg*B, χ*efg*D, and χ*efg*E exhibit distinct target cell specificities showing both functionality and altered specificity. Efagins show overlapping activity patterns, but each type displays unique activities. This indicates that each efagin targets a unique aspect of the variable and complex Rha-CWPS/Epa structure, ^42,43^ as well as more broadly shared elements. As an additional proof of targeting, χ*efg*A was used to select for resistant mutants of its prototype strain target, V583. Again, two mutants were obtained and sequenced, V583-R1 and -R2, and again both also possessed mutations in variable region genes of its *epa* operon (Extended Data Fig. 6 and Table S17). All strains targeted by chimeras were confirmed to also be targeted by filtrates of the donors from which the *ef1291* sequence originated.

### Efagin/Epa compatibility as a driver of horizontal gene exchange

Gene clusters encoding efagin (in *E. faecalis* V583, *ef1276* – *ef1293*) and Epa (*ef2198* – *ef2164*) are not linked and separated by approximately 1/3^rd^ of the genome. We never detected a population collapse associated with efagin production by an *E. faecalis* strain, nor do chimeric efagins kill the strains from which the corresponding C-terminal encoding sequence derived (Fig. 5), indicating that its ecological role is not likely fratricide.^44^ We therefore examined patterns of co-occurrence of efagin and corresponding *epa* genotypes in 493 *E. faecalis* genomes representing all major phylogenetic branches across the species (Supplementary Fig. 5). As predicted, the results reveal a non-random pattern with certain pairings occurring at high frequency (*e.g.*, efagin type B and *epa* type 8; or efagin type C and *epa* type 1), while others appear less common but still permissible. Both efagin types and *epa* genotypes are scattered across the whole genome-based phylogeny (Supplementary Fig. 6), these co-occurrences support a functional linkage and possible co-evolution between efagin and *epa*, likely shaped by selective pressures related to self-immunity.

The association between efagin type and a non-targeted *Epa* genotype in the producing cell was found to be highly significant, providing further support for a model where efagins target Epa types heterologous to that of the producer (Fisher’s exact test p < 0.00001) (Extended Data Fig. 7 and Table S19). However, exceptions were observed, both implying unexpected susceptibility as well as resistance (Extended Data Fig. 7). For example, strain SS7 (*epa* type 9) was found to be resistant to killing by efagin type C, whereas other members of *epa* type 9 were found susceptible. Close examination of the SS7 chromosome shows that it harbors an additional gene within the *epa* operon (Extended Data Fig. 7, orthogroup 151; pink), potentially modifying its Epa structure. Other exceptions without obvious gene gains or losses (*e.g.*, strain Fly2, *epa* operon type 1 but resistant to efagin type D) likely have more subtle changes such as point mutations that affect expression of specific genes within their *epa* operons, as was found in experimentally derived resistant mutants, but this remains to be shown.

### Efagins and phages select for cross-resistance

Chatterjee and colleagues^45^ observed that mutants of *E. faecalis* strain X98 gained immunity to infection by some phages after acquiring mutations in *epa* genes. We obtained two mutants (51RS1 and 51RS2) from that study to test for alteration in efagin susceptibility. Wild type X98 is targeted by efagin types A, C, and E (Extended Data Fig. 6). Phage-resistant mutant 51RS1 (with lesion in the *epa* gene EFOG_0058^45^) retains susceptibility only to types A and E, but not type C, whereas phage-resistant X98 mutant 51RS2 (with lesion in *epa* gene EFOG_0052)^45^ is resistant to all efagin types (Extended Data Fig. 6). This shows that phage selection contributes to variation in efagin susceptibility and likely vice versa.

### Efagin target range extends beyond *E. faecalis* to other enterococcal species

We previously identified structural similarities between the *epa* operons of *E. faecalis*, *E. faecium*^37^ and in the related operons of other *Enterococcus* species.^38^ It therefore was of interest to determine whether efagin activity extends beyond *E. faecalis*. A collection of 58 *E. faecium* genomes spanning the species^46^ was assessed for efagin susceptibility *E. faecium epa* operons also possess a conserved and then variable genes and assessment of their diversity was updated using currently available data (Extended Data Figs. 8 and 9; Tables S20 and S21). Of those 58 diverse strains, 35 were found to be susceptible to at least one efagin type (Extended Data Fig. 9). Interestingly, 39% of VRE *E. faecium* (7/18) are targeted by an efagin, whereas 70% (28/40) of VSE *E. faecium* are susceptible. Importantly, only 6% (1/16) of the highly successful and often VRE clonal complex 17 (CC17) lineage is targeted by an efagin. As both *E. faecalis* and *E. faecium* species co-occur in the human gut,^47^ this raises the prospect that evolving cellular surfaces resistant to efagin inhibition may have contributed to the ecological success of the VRE *E. faecium* CC17 lineage.

In addition to *E. faecium*, 40 other *Enterococcus* species spanning all four phylogenetic clades^1,5^ were evaluated for efagin susceptibility (Extended Data Fig. 10 and Table S22). Efagins were found to be effective against many species within clades I, II, and III, predominantly species possessing *epa* genes closely related to those of *E. faecalis*. In contrast, many clade IV species lack close homologs to genes for synthesis of the core rhamnopolysaccharide stem of Epa,^38,42^ and their Rha-CWPS/*epa* operon counterparts have received little study. Interestingly, *E. testudinis* was found to be susceptible to efagin despite lacking detectable sequence identity to core or variable *epa* genes of *E. faecalis* V583, suggesting the presence of a distinct cell wall polysaccharide structure that nevertheless possesses terminal decorations, or perhaps other cell wall structures, that serve as a binding target for efagin type C.

## DISCUSSION

Here we identify efagins as a new class of independently evolved, selective microbial inhibitors core to the biology of *E. faecalis*. A phage tail fiber-like protein, EF1291, targets efagins to variation in enterococcal cell wall Rha-CWPS Epa which is common to most but not all enterococcal species. Whether functional targets occur on microbes with non-rhamnose-based carbohydrate structures on their surface is currently unknown, but appears possible based on the observed inhibition of *E. testudinis*. With activity against *E. faecium* strains, including some VRE clinical isolates, and the demonstrated ability to engineer specificity through discrete domain swaps, such agents and potentially engineered derivatives hold promise for selectively controlling infection as well as the gut ecology of hospitalized patients limiting transmission and spread.

The discovery of efagins provides a new perspective on *E. faecalis* ecology and evolution as well. Their conservation in all *E. faecalis* and their uniqueness to that species imply that they were either gained at the time of species radiation over 300 million years ago^1^ and transmitted vertically, or later acquired and spread throughout a population sweep. Their invariant occurrence in widely ranging ecologies, and absence outside of that species, argue strongly for the former. With activity against competing strains of many enterococcal species, efagins appear well suited to facilitate *E. faecalis* entry into diverse gut consortia. That is, as animals along with their microbes are consumed up the food chain, *E. faecalis* in prey may deploy efagins in the gut of a predator to displace competitors. As facile gene exchangers^48^ the escalation of *E. faecalis* up the food chain from a vast range of ecosystems positions them ideally to be important conduits of adaptive traits from various ecosystems, including naturally occurring antibiotic resistance genes.

Cross selection and coevolution of efagins, phages and their Epa targets has likely contributed to *E. faecalis* genetic diversity. Here, we observed that the evolutionary dynamics of the efagin and *epa* differ from much of the rest of the core genome, raising the prospect that efagin-mediated selection may represent a new important driver of gene exchange. Efagin susceptibility patterns across varying *epa* operon genotypes likely mirror phage susceptibility patterns, substantially informing the development of phages for therapeutic purposes as well.

The discovery of efagins provides yet another independently derived example of what is often viewed as the co-option and redeployment of a phage structure for the selective advantage of a bacterium.^49^ However, as for other phage component-like bacterial factors, the paucity of intermediates makes it difficult to substantiate a precise evolutionary pathway, making it entirely possible if not likely that the independent utility of tail-like (or alternatively capsid-like) functional units has simply been co-opted in parallel by microbes and viruses for various cellular and viral uses. ^50^

As for tailocins, ^15,18,21,24,49,51^ we show that efagins are capable of inhibiting the growth of targeted strains under the conditions tested here. However, much remains to be learned about the actual ecological role of efagins in nature. The precise mechanism of action for efagins, as for other non-contractile F-type tailocins, is unclear^18,21,22^. If competitor killing or inhibition is the objective, the nature of the selection that favors a 160 nm tail polymer is also unclear, when a) shorter variations spontaneously occur, and b) killing involves activities mainly of the baseplate.^22^

Much remains to be learned about the contributions of efagins to the biology of *E. faecalis*, and their contribution to its exceptional success – both as a commensal of a wide range of terrestrial animal guts and as a persistent, multidrug resistant hospital pathogen. Irrespective of their role in nature, they clearly possess the ability to be engineered to target cells expressing a cognate cell wall rhamnopolysaccharide (and possibly related structures), which may be of considerable value in managing and limiting the spread of multidrug resistant enterococci in medical and agricultural environments. In an era when few effective treatment options remain for those with multidrug resistant enterococcal infections, the identification of a new agent that targets these recalcitrant organisms represents an important advance.

## METHODS

### Collection of bacterial strains

The bacterial collection used in the experimental work is described in Tables S23, S24, S25, and S26. *Enterococcus* species were propagated in Brain Heart Infusion (BHI) broth or on 1.5% BHI agar at 34 °C or 37 °C, depending on temperature preferences for each species^5^, while *Vagococcus giribetii* was incubated at 34 °C. *Escherichia coli* was maintained on 2x Yeast Tryptone medium (2xYT) at 37 °C.

### Induction and recovery of efagin particles

To obtain efagin particles, bacterial cultures were grown overnight in BHI broth and incubated at 37 °C for 18 h. For strains not expressing efagin constitutively, overnight cultures were diluted 1:100 in BHI broth and incubated for 4 h, followed by induction with mitomycin C (final concentration 2.5 µg/ml) and incubation at 37 °C overnight. The cell debris was removed by centrifugation at 11,000 × *g* for 15 minutes at 4 °C. Then the supernatant was collected and filtered through a 0.2 um membrane (Pall Life Sciences). Efagin particles collected in the supernatants were added along with 1 M NaCl (5.84 g/100 ml) and 10% (w/vol) polyethylene glycol 8000 (PEG), and the mixture was incubated for 1h at 4 °C. The PEG-precipitated material was then collected by centrifugation at 25,000 × *g,* 4 °C, for 1 h. The pellet was resuspended in SM buffer (50mM Tris-Cl pH 7.5, 100 mM NaCl, 8mM MgSO_4_•7H_2_O,) and another addition of 2.5 M NaCl and 20% (w/v) polyethylene glycol PEG-8000 was performed for 30 min at room temperature. Efagin particles were collected by centrifugation at 17,000 × g, 4 °C, for 1 h, and the pellets were resuspended in SM buffer.

### Efagin purification and concentration

Efagin particles were purified and concentrated using centrifugal filter units: Macrosep 100 K MWCO advance centrifugal devices (20 ml; Pall Life Sciences), and Amicon Ultra 100K; (0.5 ml; Millipore) according to the manufacturer’s instructions. Briefly, efagin suspensions in SM buffer were loaded into Macrosep 100 K and centrifuged at 4,000 × g for 1.5 h. The collection tubes were then emptied, refilled with 10 mL SM buffer, and subjected to a second centrifugation step. Retained particles were resuspended in 2 mL SM buffer and transferred to Amicon Ultra 100 K filters. These were centrifuged at 5,000 × g for 30 min, and efagin was recovered by reverse spinning at 1,000 × g for 2 min. The resulting concentrated preparations were used for activity assays of efagin and for subsequent analyses.

### Transmission Electron Microscopic (TEM)

Purified efagin particles were diluted in SM buffer to a concentration of 10^7^ efagin-forming units/ml. Freshly ionized carbon coated grids were floated on a 10 ul drop of sample for 1 minute. Grids were washed with 5 drops of 2% uranyl acetate (UA), and the excess of UA was drawn off with grade 50 Whatman filter paper. Grids were allowed to dry and imaged with a Hitachi 7800 at 100KV.

### Grids preparation and imaging conditions for single particle analysis

Purified efagin particles in a volume of 3 µl were applied onto the glow discharged Quantifoil R 1.2/1.3, 300 Mesh, Au grid and plunge frozen using the Vitrobot Mark 4 (Thermo Fisher Scientific, Waltham, MA, USA) in 100% humidity and 4°C. The grids were blotted for 3s and automatically plunged into liquid ethane. Vitrified samples were transferred to storage boxes and stored in liquid nitrogen until use. Grids were clipped in AutoGrid Rings and loaded into a 300 kV Titan Krios electron microscope (Thermo Fisher Scientific, Waltham, MA, USA) with Gatan K3 Camera and BioQuantum Energy Filter set to a 20 eV slit width. Micrographs were collected with a nominal 81.000x magnification, with a corresponding pixel size of 1.065Å, a total dose equal 60e-/Å, 50 frames and defocus range of -2.1 to -0.9 µm.

Micrographs were imported to the CryoSPARC v4.6 software,^52^ followed by Patch Motion Correction and Patch CTF Correction. Three hundred random efagin particles were manually measured, which allowed us to prepare the length distribution plot using RStudio v2025.09.2 software.^53^

### Detection of phage activity and titration

Efagin particles from OG1RF *E. faecalis* were tested on an indicator strain for plaque formation.^54^ Briefly, the indicator strain was cultured overnight and then diluted to equal the turbidity of a 0.5 McFarland standard (∼1.5 x 10^8^ colony forming units (CFU)/ml). One hundred microliters of the bacterial suspension were mixed with 1 ul of diluted efagin particles (prepared 10 to 10^9^-fold serial dilutions) and 5 ml of 0.5% BHI agar, which was then poured onto a 1% BHI agar plate. Once the overlay solidified, the plates were air-dried and incubated overnight at 37 °C. Plaque formation was visually detected and enumerated by plaque assay using JM110 *Escherichia coli* (indicator strain) and M13 bacteriophages as the positive control.

### Efagin locus knockout in *E. faecalis* OG1RF

An efagin knockout strain of *E. faecalis* OG1RF was generated through allelic replacement. Genomic regions flanking the efagin locus (878 bp upstream and 860 bp downstream) were amplified by PCR using primer pairs KO-F1_XbaI up-forward/KO-R1_Acc65I up-reverse and KO-F2_KpnI down-forward/KO-R2_SbfI down-reverse (Table S27), then fused via overlap extension PCR to create a 1,738 bp deletion construct. This fragment was digested with *XbaI*/*SbfI* and cloned into the temperature-sensitive plasmid pLT06 pre-digested with the same enzymes. The recombinant plasmid was propagated in *E. coli* EC1000 and verified by sequencing. Following PCR confirmation with primers OriF and KS05seqR, the construct was introduced into *E. faecalis* OG1RF by electroporation. Allelic exchange was carried out using temperature shift and p-chlorophenylalanine counterselection, as described previously.^41^ Candidate mutants were screened by PCR using internal primers (efagin-internal-F and efagin-internal-R; **Table S27**), and successful deletion was confirmed by whole genome sequencing.

### Construction of efagin chimeras in *E. faecalis* OG1RF

Efagin chimeras of *E. faecalis* OG1RF were generated by allelic exchange. A 1,324 bp DNA fragment spanning the region between the C-terminal end of *ef*1291 and the start of the EF1292 efagin gene was amplified by PCR using primer pair 817_F_BamHI up-forward/2150_R_PstI up-reverse (Table S27). The resulting product was digested with BamHI and PstI and ligated into the temperature-sensitive plasmid pLT06, pre-digested with the same enzymes. Recombinant plasmids were propagated in *E. coli* EC1000 in medium supplemented with chloramphenicol (15 μg/mL), verified by PCR using primers OriF and KS05seqR, and confirmed by Sanger sequencing. Plasmids were subsequently extracted and introduced into *E. faecalis* OG1RF by electroporation. Allelic exchange was carried out using temperature shift and p-chlorophenylalanine counterselection, as previously described.^41^ Candidate mutants were screened by PCR using internal primers for efagin types A–E (internal-F and internal-R; Table S27) and further evaluated by activity assays. Successful recombinants were validated by sequencing.

### Host range determination

Efagin killing assays were conducted using spot tests and the double agar layer method. ^55^ The indicator strains were cultured overnight and then diluted to equal the turbidity of a 0.5 McFarland standard (∼1.5 x 10^8^ CFU/ml). One hundred microliters of the bacterial suspension were applied to a 0.5% Brain Heart Infusion (BHI) agar overlay, which was then poured onto a 1% BHI agar plate. Once the overlay solidified, the plates were air-dried and incubated in the refrigerator for 1 hour. Following this, 4 μl of efagin preparations were carefully spotted onto the plates and allowed to dry/adsorb and incubated at 4°C for 2h. After that, the plates were incubated at 37 °C overnight, and zones of clearing indicated positive bactericidal activity.

### Isolation of efagin-resistant *E. faecalis* strains

The efagin-resistant phenotypes were selected using spot tests and the double agar layer method as described previously. Plates were let dry, and 4-20 µl of efagin particles were spotted and allowed to dry/adsorb for 4 h. After, plates were incubated at 37 °C overnight. The presumptive efagin-resistant colonies appeared in the zones of clearing. The efagin-resistant phenotypes were confirmed by spot assays and whole genome sequencing (WGS).

### Genomic DNA preparation, high-Throughput sequencing, assembly, and annotation

*Enterococcus* strains were cultured in BHI broth at 37 °C for 18 h. Genomic DNA was extracted using the DNeasy Blood & Tissue Kit (QIAGEN, San Luis, MO, USA) with minor modifications to the manufacturer’s protocol. Specifically, 50 μL of lysozyme (50 mg/mL) and 10 μL of mutanolysin (2,500 U/mL; Sigma-Aldrich, Germantown, MD, USA) were added and incubated for 30 min at 37 °C prior to treatment with 20 μL proteinase K (20 mg/mL). DNA concentrations were determined using the Qubit dsDNA High-Sensitivity Assay Kit (Thermo Fisher Scientific, MA, USA). Sequencing libraries were prepared with the Nextera XT DNA kit and Illumina index primers. Paired-end reads (2 × 250 bp) were generated on an Illumina MiSeq platform with the MiSeq Reagent Kit v2. Reads were trimmed to remove adapters, quality-filtered, and screened for PhiX contamination before de novo assembly in CLC Genomics Workbench v8.0.3. Ten additional strains (efagin resistant-mutants and chimeras) were sequenced externally by CD Genomics (https://www.cd-genomics.com) and processed using the same pipeline.

All Illumina sequencing reads and assembled genomes have been deposited in GenBank under BioProjects PRJNA313452^56^ and PRJNA663900,^57^ and PRJNA1371623^58^. Strains and corresponding accession numbers are provided in Table S1.

### WGS variant calling analysis

Whole-genome sequencing was performed on both efagin-resistant mutants and their corresponding parental strains to identify single nucleotide polymorphisms (SNPs) and insertions/deletions (indels). Sequencing reads were aligned to the finished reference genome using the BWA-MEM algorithm (v0.7.4).^59^ Variant calling was then carried out using Pilon (v1.23).^60^ Variants with a mapping quality score below 10 were excluded from the final dataset to ensure accuracy.

### Searching for efagin sequence in Enterococcaceae and other prokaryotic bacteria

We downloaded all 16,907 Enterococcaceae assemblies (Taxid 18,015) from NCBI (https://www.ncbi.nlm.nih.gov), in May 2021, to analyze together with additional Enterococcaceae genomes from the Broad Institute’s collection (n= 1108). Quality controls were performed to remove contaminated or incomplete genomes, using Phylosift^61^ and CheckM.^62^ FastANI^63^ was used to identify species in our dataset by comparing to reference genomes, including *E. faecalis* V583, with the presence of >95% average nucleotide identity to a known reference genome confirming the species designation. Our final dataset contained 15,854 confirmed Enterococcaceae assemblies for analysis, including 3,068 *E. faecalis*.

BLAST (https://www.ncbi.nlm.nih.gov) was used to search our enterococcal assembly database for hits matching the efagin sequence from *E. faecalis* reference strain V583. We calculated the total coverage of efagin for hits within each target sequence, accounting for overlapping hits. Efagin was defined as present in a strain if there were Blast hits with at least 90% coverage of efagin (E-value <10), including at least one hit of length > 2000 bp. Another, more comprehensive BLAST search was performed against all draft prokaryotic genomes available on NCBI (https://www.ncbi.nlm.nih.gov). All sequencing submissions matching the query “wgs_master[prop] AND (bacteria[orgn] OR archaea[orgn]) NOT refseq[filter]” on the NCBI Nucleotide website were included in this dataset (https://www.ncbi.nlm.nih.gov), or approximately 1.2 million sequencing submissions in Oct 2022.

### Selection of representative *E. faecalis, L. monocytogenes, P. aeruginosa and C. difficile* genomes to cover the breadth of diversity within the species

In order to remove biases within our dataset due to over-representation of commonly sequenced clinical subtypes, and to ensure computational tractability, we selected representative genomes for each species, chosen to represent the diversity present across the species. Briefly, all genomes were clustered using the StrainGR (v1.3) software package^64^, using its utility for creating representative reference genome databases. We clustered genomes with a jaccard distance of 0.9, and a representative was chosen for each cluster.

For *E. faecalis*, we started with our full set of 3,068 genomes. After clustering, we chose the representatives with the smallest number of contigs, that also contained a full efagin on a single contig, resulting in a set of 441 *E. faecalis* genomes. Additionally, we added lab-tested strains important to this study, with a final total of 493 genomes representing the diversity of *E. faecalis*.

For analyses of *Listeria monocytogenes*, *Pseudomonas aeruginosa*, and *Clostridium difficile*, we started with 414, 883 and 189 genomes from the complete refseq database, respectively. After clustering, the resulting datasets contained 74, 343, and 32 genomes, respectively, highlighting different levels of diversity for each species in the database.

### Searching for partial phages

To detect partial phages from additional enterococcal species and closely related *Melissococcus plutonius*, *Tetragenococcus halophilus*, and *Vagococcus fluvialis*, we used all isolates for each species from the refined dataset of 15,854 Enterococcaceae assemblies described above. These were then used as input into the provirus identification tool within the geNomad software package. ^10^ This tool identifies and extracts proviruses from assembled genomes by detecting regions enriched in viral hallmark genes using genomad’s neural network classification model. We filtered for proviruses present in the majority of isolates in each species and explored whether they lack typical prophage proteins using geNomad^10^. Incomplete annotations were confirmed with Phastest.^11^

### Identification of the closest known ancestor of efagin and protein identity

All V583 efagin gene sequences were analyzed using the phage prediction tool PHASTER^65^ and Phastest^11^ to identify the closest known phage ancestor. The resulting phage protein sequences were then compared with efagin proteins using BLAST alignments (default E-value thresholds) to determine protein-level identity.

### Monocin similarity analysis

Using nucleotide sequences from all V583 efagin genes, we searched against the monocin gene sequences and obtained no hits (default E-value thresholds). Using sequences from all V583 efagin proteins, we searched against the monocin protein sequences from ATCC 35152 strain using a BLAST e-value threshold of 1e-5.

### Functional annotation of efagin and genes in its genomic neighborhood

Gene function information for efagin (*ef1276* to *ef1293*) and its surrounding region (20 genes upstream and downstream) was obtained by combining information from Prokka,^66^ Pfam,^67^ Kegg,^68^ Phyre 2.2,^29^ BioCyc,^69^ Alphafold 3,^31^ and PaperBlast.^70^ The annotated genes immediately upstream and downstream of *efagin* via this pipeline are not present in the *E. faecalis* V583 genome annotation in GenBank (https://www.ncbi.nlm.nih.gov), but were found in our Prokka^66^ annotations. For reference in our analysis, we assigned them arbitrary *ef* IDs: *ef3441* (upstream) and *ef3442* (downstream).

### Orthogroup clustering and characterization of *epa*

To obtain efagin particles, bacterial cultures were grown. Among all isolates containing a complete efagin sequence, 1,972 also had a complete epa operon on one contig. Since *epa* consists of a conserved region followed by a variable region (containing 7 to 27 genes), a conserved gene downstream of *epa*, *ef2162*, was used to define the downstream boundary of the operon. The genes between *epa*A and *ef2162*, not including *ef2162*, were considered part of the *epa* operon and were included in our orthogroup clustering analysis. SynerClust^71^ was run on these genomes, classifying all *epa* genes into 137 orthogroups. To further correct for possible overclustering artifacts, we split orthogroups with within-cluster average nucleotide identity <90%.

To explore the presence of *epa* in *E. faecium*, we downloaded 58 diverse *E. faecium* genomes from a previously published study^46^ and repeated the same orthogroup clustering process described above. In total, 95 epa orthogroups were present in *E. faecium*.

In order to understand epa diversity across the *Enterococcus* genus, we used previously published results^38^ that used BLAST to identify which epa genes are present in other enterococcal species, calculating the percent identity between genes from different enterococci and the *E. faecalis* V583 epa genes. For new isolates not included in that study,^38^ we also used the same method to detect epa genes, using the same E-value threshold of 1E-10 for inclusion as a positive hit.

### Analysis of correlation between efagin SNPs and *epa* genes

For each gene in the variable region of the epa, we created a vector with a binary value corresponding to the presence or absence of this gene within each *E. faecalis* representative genome selected as described above. Correspondingly, for each 150bp region within efagin, we created a vector containing the number of SNPs in that region for each representative genome within our database. A position in a genome was considered a SNP if it differed from the nucleotide at that position in V583 or if it was an inserted or deleted relative to V583. Pearson correlations were calculated between the vectors for each epa gene and each efagin region.

### Protein structural modeling

Alphafold 3^31^ was used to model the full-length protein structures of all five efagin *ef*1291 types via alphafoldserver.com. As gp15^30^ is known to form a homotrimeric structure, we included three copies of *ef*1291 as input. We generated visualizations using Pymol.^72^

### Defining Epa and efagin types

We defined epa types using hierarchical clustering, based on the presence or absence of *epa* genes in our orthogroup clustering based on 1,972 *E. faecalis* genomes. This was performed using the AgglomerativeClustering package from scikit-learn.^73^ We chose an optimal threshold by comparing the clusters with patterns of efagin killing activity, achieving optimal agreement with the distance_threshold parameter value of 9. For the 58 *E. faecium* genomes, we applied the same approach, using a distance_threshold parameter value of 6.

Efagin types were defined for the complete dataset using hierarchical clustering of an alignment of amino acid residues representing the variable region of *ef*1291 (V583 efagin coordinates 12,539-12,967). We used the AgglomerativeClustering package,^73^ and an optimal threshold was chosen by comparing the clusters with patterns of efagin killing activity, achieving optimal agreement with a distance_threshold value of 24.

### Construction of phylogenies

The whole-genome phylogeny of *E. faecali*s representatives was generated with Parsnp v1.1.2 ^2^/23/26 4:23:00 PM using default parameters and V583 as the reference genome. To construct the single-copy core phylogenetic tree of Enterobacteriaceae, individual representative genomes for each species were used as input to Orthofinder^74^ to find all genes present in >=95% of isolates. These were aligned with MAFFT^75^ and then phylogenetically reconstructed using IQTree^76^ in modelfinder mode with 1000 ultrafast bootstraps.

## Data availability

All Illumina sequencing reads and assembled genomes have been deposited in GenBank under accession numbers BioProjects PRJNA313452^56^, PRJNA663900^57^, and PRJNA1371623^58^. Strains and corresponding accession numbers are provided in Tables S1. Any additional information required to reanalyze the data reported in this paper is available from the lead contact (Michael S. Gilmore) upon request.

## Supporting information

Graphical Abstract

Highlights

## ACKNOWLEDGMENTS

We thank Nicki E. Watson and the Harvard Center for Nanoscale Systems for technical assistance with TEM. This work was supported by NIH grant AI083214 (Harvard-wide Program on Antibiotic Resistance) and was also funded in part with federal funds from the National Institute of Allergy and Infectious Diseases, National Institutes of Health, Department of Health and Human Services, under grant numbers U19AI110818 (Broad Institute) and F32AI179151 (J.T.S). Work in the Kaplan laboratory at UChicago was supported by the National Institute of General Medical Sciences (R35GM157116 to M.K.) and the Searle Scholars Program (to M.K.). We gratefully acknowledge the expert technical assistance and support provided by Ryan Whipple and Jose T. Saavedra. The authors also thank members of the Gilmore and Earl Labs for helpful discussions during the development of this project and preparation of the manuscript.

## AUTHOR CONTRIBUTIONS

M.S.G. conceived the study. M.S.G. and J.P. designed the overall experimental work and A.M.E. and A.L.M. supported project development and provided guidance on both the computational analyses and the biological aspects. A.L.M., J.T.S., G.R., and J.P. performed all the bioinformatic analyses. N.K. conducted the initial detection of efagin activity, and the efagin deletion mutant in OG1RF was constructed by N.K. and S.S.M. J.P. optimized efagin production and purification, performed activity assays, and generated efagin-resistant mutants and efagin chimeras. A.C. and M.K. performed the efagin particle measurements. J.P., A.L.M., J.T.S., A.M.E., and M.S.G. contributed to data interpretation and scientific discussion. J.P. and M.S.G. wrote the original draft, and J.P., A.L.M., J.T.S., and M.S.G. revised and edited the manuscript. A.M.E. and M.S.G. acquired funding and supervised the project. All authors read and approved the final manuscript.

## ADDITIONAL RESOURCES

All genomes used in this study are described in **Tables S1, S5, S7, S11, S24, and S25.**

Broad Institute. Data from Bioproject PRJNA313452. NCBI Genbank. https://www.ncbi.nlm.nih.gov/bioproject/?term=PRJNA313452.

Broad Institute. Data from Bioproject PRJNA663900. NCBI Genbank. https://www.ncbi.nlm.nih.gov/bioproject/?term=PRJNA663900.

Mass Eye and Ear/Harvard Medical School. Data from Bioproject PRJNA1371623. NCBI Genbank. https://dataview.ncbi.nlm.nih.gov/object/PRJNA1371623.

## EXTENDED DATA FIGURES

**Extended Data Fig. 1. Genomic neighborhood of efagin shows conserved flanking genes**. Bar plots display the frequency of genes found upstream (left) and downstream (right) of efagin across *E. faecalis* genomes. The most common gene at each position is shown in red, the second in green, and the third in purple.

**Extended Data Fig. 2. Comparison of gene clusters encoding phage tail-like elements.** Pyocin (30.3 kb), diffocin (21.8 kb), monocin (11.6 kb), and efagin (14.6 kb) from *Pseudomonas aeruginosa*, *Clostridium difficile*, *Listeria monocytogenes*, and *Enterococcus faecalis*, respectively. While these systems show broad similarity in gene organization, they share little to no sequence identity, suggesting independent evolutionary origins. Genomic comparison between the *L. monocytogenes* monocin locus and the *E. faecali*s efagin locus reveals similar gene organization, particularly in structural modules such as the tube, tape measure, baseplate components, and receptor-binding protein (RBP). Although efagin shares organizational similarity with monocin, it has minimal sequence identity with monocin across three genes (red asterisk). Both systems encode non-contractile phage tail-like structures and appear to have arisen independently. Arrows represent open reading frames (ORFs), color-coded by predicted function: regulatory (lightly grey), structural (pink), and lysis-associated genes (purple). Particles are classified as R-type (contractile) or F-type (non-contractile). Reference sequences used include: i) the R- and F-type pyocin clusters from *Pseudomonas aeruginosa* PAO1, corresponding to the genomic region spanning genes PA0610–PA0648 (Genbank: AE004091.2); ii) the R-type diffocin from *Clostridium difficile* 630 (GenBank: NZ_CP010905.2; genes CD1359–CD1379); iii) the F-type monocin from *Listeria monocytogenes* 10403S (GenBank: NC_017544.1; genes LMRG02361–LMRG02378); and iv) efagin from *E. faecalis* V583 (GenBank: NC_004668.1; genes ef1276–ef1293) were included as reference sequences. The number inside each arrow denotes the number of genes represented.

**Extended Data Fig. 3. Distribution of contractile (R-type) and non-contractile (F-type) phage tail-like elements among genomes from Gram-positive and Gram-negative bacterial species.** In contrast to efagin, which is highly conserved and intact in all strains for which high-quality genome sequences are available, the other tailocins appear variable and often occur in alternate hybrid forms.

**Extended Data Fig. 4. Predicted model for EF1291 as ribbon and space-filling structures using Alphafold 3.**^31^ The three colors (blue, red, and yellow) represent individual monomers within the trimer. Regions highlighted with darker colors in the C-terminal end correspond to the EF1291 structural regions related to the receptor-binding domain of Listeria phage PSA protein gp15 by Phyre2.2.^29^

**Extended Data Fig. 5. Coordinated variation of *epa* gene presence/absence and efagin-associated SNPs across *E. faecalis* representative genomes.** Example of a correlation between a sequence variant in efagin *ef*1291 and the presence or absence of a specific gene in the *epa,* across the diversity of *E. faecalis* strains. This plot shows variation in position 12903 of *ef*1291 and the presence or absence of the *epa* gene *ef*2167. Gray shading indicates presence or absence concordance between mutated position 12903 and *ef*2167. Similar correlations were observed between efagin positions and *ef*2166, *ef*2167, *ef*2168, and *ef*2169, four adjacent genes within the *epa*.

**Extended Data Fig. 6. Efagin resistant mutants have defects in *epa* variable region genes. a,** Effect of *epa* mutations on efagin susceptibility in four *E. faecalis* strains (FA2-2, X98, V583, and OG1RF). **b,** Other studies^43,45,78^ describing *epa* mutations or indels correlated with phage resistance

**Extended Data Fig. 7. Relationship between efagin activity and *epa* operon architecture across 54 *E. faecalis* strains.** Schematic figure showing the left-hand heat maps display efagin activity assays from five different donor strain: OG1RF (type C efagin) and OG1RF chimeras: *χefgA* (type A efagin), *χefgB* (type B efagin), *χefgD* (type D efagin), and *χefgE* (type E efagin), annotated with its *epa* operon type and assigned efagin type (A-E) against a diverse collection of *E. faecalis* strains (n=54). Grey indicates susceptible and white non-susceptibility. Grey hashing highlights donor–target pairs that share either the same efagin (hashes pointing up to the right) or/and *epa* type (hashes pointing up to the left), indicating situations where we would not expect inhibition, and most often do not observe inhibition (**Table S19**). Operon cluster IDs are listed for each target strain. The figure highlights how variations in *epa* gene content, and efagin type are associated with differential killing profiles. Chimeric constructs are denoted using the Greek letter “*χ*” (chi), and efagin is abbreviated as “*efg*.”

**Extended Data Fig. 8. Comparison of the organization of the *epa* locus between *E. faecalis* V583 and *E. faecium* Aus0004.** The epa rhamnose backbone is conserved in both species (dark yellow arrows), but differs in gene composition and arrangement, whereas the variable region (light yellow arrows) is completely different in the two species. Core epa genes (epaA–epaR) show moderate to high sequence identity (31.2–92.1%), whereas the variable decoration genes (light yellow) lack detectable homology, highlighting species-specific diversification of surface polysaccharide structures. In E. faecium, the locus follows the order epaABCDEFGHPQLMNOR. The E. faecium epaN gene resembles E. faecalis epaN at the N-terminus, sharing a predicted S-adenosylmethionine binding site, but differs in its C-terminal sequence. We identified orthologs of epaP and epaQ in both species, but located at distinct genomic sites. In E. faecalis, the central region of the cluster contains epaI, epaJ, and epaK, whereas in E. faecium these genes are absent and replaced by epaP and epaQ, suggesting the rhamnose polysaccharides of the two species have different structure.

**Extended Data Fig. 9. Structural diversity of *epa* operons across *E. faecium* strains and correlation with efagin susceptibility**. Strain identifier, sequence type (ST), clonal complex (CC), *E. faecium* clade assignments (A1, A2, or B) and vancomycin resistance genotype in relation to the organization of *epa* operons. Rec indicates strains that are hybrids between clades. Conserved core genes are shown in black, while variable genes are colored according to orthogroup membership, with shapes corresponding to predicted function. The rightmost panel shows the efagin susceptibility profile. In the Epa cluster, black represents core genes found in all the genomes, and grey indicates rare genes found in less than 5%.

**Extended Data Fig. 10. Conservation of *epa* operon genes across *Enterococcus* species and correlation with efagin susceptibility**. Strains of *Enterococcus* clades are colored in graded shades of purple by membership in clades I – IV.^1^ The extent of amino acid sequence identity with inferred *Epa* gene products, when compared to the prototypes of *E. faecalis* strain V583, is indicated by the gold heatmap. Efagin susceptibility is indicated with grey shading.

## SUPPLEMENTARY FIGURES

**Supplementary Fig. 1. V583 efagin gene function prediction**. Biocyc, PaperBlast, Phyre2.2, Blastp, Prokka, Pfam, and Alphafold were used to infer the function.

**Supplementary Fig. 2. Variation of *epa* gene presence/absence and efagin-associated SNPs across *E. faecalis* representative genomes.** This plot shows the distribution of *epa* gene presence (yellow) and efagin *ef*1291 SNPs (red) across *E. faecalis* strains, aligned to a whole-genome phylogeny (center). Concordance between these *ef*1291 SNPs and the presence of these specific *epa* genes was observed in 81% of strains, far exceeding the ∼50% expected by chance, suggesting coordinated evolutionary dynamics. The enrichment of these reciprocal patterns supports a functional or selective relationship between efagin and the *epa* locus.

**Supplementary Fig. 3. Visualization of efagin from OG1RF*χefgD* chimera strain.** Transmission electron microscopy (TEM) shows images of efagin particles produced by the *E. faecalis* OG1RF*χefgD* chimera strain expressing type D efagin. Morphological features were visualized and quantified using TEM and analyzed with Fiji/ImageJ software.

**Supplementary Fig. 4. Gene structure of representatives of *Epa* type 11 in 10 *E. faecalis* genomes**. Each row represents a distinct *E. faecalis* genome, labeled by its NCBI accession number. Conserved core *epa* genes are shown in black, while variable genes (e.g., 000192, 000190, etc.) are represented in distinct colors, highlighting differences in the epa operon across strains, within *Epa* type 11.

**Supplementary Fig. 5. Frequency of combinations of efagin and EPA types present in 493 *E. faecalis* representative genomes.** Rows represent different efagin types; columns represent different *epa* types. The central heat map indicates the frequency of each combination of *epa* and efagin types; marginal bar plots show the frequency of each *epa* or efagin type individually across the 493 genomes. A white square indicates that there are no genomes with that combination of efagin and *epa* types.

**Supplementary Fig. 6. Whole-genome phylogeny of *E. faecalis* strains, overlaid with the diversity of *epa* types (outer ring; 1-20) and efagin types (inner ring; A-E, and X).** The lack of concordance between the genomic phylogeny and the distribution of efagin or epa types suggests that efagin and *epa* genetic diversity were likely acquired by horizontal gene transfer and localized recombination. Scale bar indicates genetic distance.

## SUPPLEMENTARY TABLES

**Table S1.** BLAST results of efagin against 3,068 high-quality *E. faecalis* genomes

**Table S2.** Summary of Genomad-identified phage-derived elements in *Melissococcus plutonius*

**Table S3.** Summary of Phastest-identified phage-derived elements in *Melissococcus plutonius*

**Table S4.** Results of BLAST analysis comparing efagin and monocin

**Table S5.** Summary of monocin loci identified in 74 *Listeria monocytogenes* genomes

**Table S6.** Monocin deletion identified across 13 *Listeria monocytogenes* genomes

**Table S7.** Summary of R-type pyocin loci detected across 343 *Pseudomonas aeruginosa* genomes

**Table S8**. R-pyocin variant region identified across 187 *Pseudomonas aeruginosa* genomes

**Table S9.** Summary of F-type pyocin loci detected across 343 *Pseudomonas aeruginosa* genomes

**Table S10.** F-pyocin variant region identified across 55 *Pseudomonas aeruginosa* genomes

**Table S11.** Summary of R-type diffocin loci detected across 32 *Clostridium difficile* genomes

**Table S12.** R-diffocin variant region identified across 18 *Clostridium difficile* genomes

**Table S13.** BLAST results for the *E. faecalis* phage most closely related to the efagin

**Table S14.** List of efagin and *epa* types assigned to 1,972 *E. faecalis* genomes included in the study

**Table S15.** BLAST results of type E efagin C-terminal end

**Table S16.** V583 efagin gene function prediction performed by Prokka, Pfam, and KEGG

**Table S17.** Whole-genome sequencing data of spontaneous efagin-resistant *E. faecalis* mutants

**Table S18.** EPA orthogroup dataset of 1,972 *E. faecalis* genomes

**Table S19.** Statistical analysis of the association between efagin or *epa* type matching and susceptibility to killing

**Table S20.** EPA orthogroup dataset of 58 *E. faecium* genomes

**Table S21.** Amino acid identity between epa core genes of *E. faecalis* V583 and *E. faecium* Aus0004 based on BLASTP alignments

**Table S22.** Amino acid sequence identity (%) between genes from the epa cluster of *E. faecalis* V583, and their top hit in 40 *Enterococcus* species

**Table S23.** List of *E. faecalis* strains used in the experimental work

**Table S24.** List of *E. faecium* strains used in the experimental work

**Table S25.** List of *Enterococcus* species used in the experimental work

**Table S26.** List of strains used in this study

**Table S27.** List of primers used in this study

## REFERENCES

1. Lebreton, F. et al. Tracing the Enterococci from Paleozoic Origins to the Hospital. Cell 169, 849–861.e13 (2017).

2. Centers for Disease Control and Prevention (CDC). Antibiotic Resistance Threats in the United States. https://www.cdc.gov/antimicrobial-resistance/data-research/threats/index.html.

3. Ikuta, K. S. et al. Global mortality associated with 33 bacterial pathogens in 2019: a systematic analysis for the Global Burden of Disease Study 2019. The Lancet 400, 2221–2248 (2022).

4. European Centre for Disease Prevention and Control - ECDC (2024). Antimicrobial Resistance in the EU/EEA (EARS-Net)—Annual Epidemiological Report 2023. November 18, 2024. https://www.ecdc.europa.eu/en/publications-data/antimicrobial-resistance-eueea-ears-net-annual-epidemiological-report-2023.

5. Schwartzman, J. A., et al. Global diversity of enterococci and description of 18 previously unknown species. Proc. Natl. Acad. Sci. 121, e2310852121 (2024).

6. Paulsen, I. T. Role of Mobile DNA in the Evolution of Vancomycin-Resistant Enterococcus faecalis. Science 299, 2071–2074 (2003).

7. Matos, R. C., et al. *Enterococcus faecalis* Prophage Dynamics and Contributions to Pathogenic Traits. PLoS Genet. 9, e1003539 (2013).

8. Scholl, D. Phage Tail–Like Bacteriocins. Annu. Rev. Virol. 4, 453–467 (2017).

9. Backman, T., Burbano, H. A. & Karasov, T. L. Tradeoffs and constraints on the evolution of tailocins. Trends Microbiol. 32, 1084–1095 (2024).

10. Camargo, A. P. et al. Identification of mobile genetic elements with geNomad. Nat. Biotechnol. 42, 1303–1312 (2024).

11. Wishart, D. S. et al. PHASTEST: faster than PHASTER, better than PHAST. Nucleic Acids Res. 51, W443–W450 (2023).

12. Bobay, L.-M., Touchon, M. & Rocha, E. P. C. Pervasive domestication of defective prophages by bacteria. Proc. Natl. Acad. Sci. 111, 12127–12132 (2014).

13. Kageyama, M., Ikeda, K. & Egami, F. Studies of a Pyocin: III. Biological Properties of the Pyocin. J. Biochem. (Tokyo) 55, 59–64 (1964).

14. Heo, Y.-J., Chung, I.-Y., Choi, K. B. & Cho, Y.-H. R-type pyocin is required for competitive growth advantage between Pseudomonas aeruginosa strains. J. Microbiol. Biotechnol. 17, 180–185 (2007).

15. Ge, P. et al. Action of a minimal contractile bactericidal nanomachine. Nature 580, 658–662 (2020).

16. Carim, S. et al. Systematic discovery of pseudomonad genetic factors involved in sensitivity to tailocins. ISME J. 15, 2289–2305 (2021).

17. Blasco, L. et al. Study of 32 new phage tail-like bacteriocins (pyocins) from a clinical collection of *Pseudomonas aeruginosa* and of their potential use as typing markers and antimicrobial agents. Sci. Rep. 13, 117 (2023).

18. Saha, S. et al. F-Type Pyocins Are Diverse Noncontractile Phage Tail-Like Weapons for Killing *Pseudomonas aeruginosa*. J. Bacteriol. 205, e00029–23 (2023).

19. Zink, R., Loessner, M. J. & Scherer, S. Charaterization of cryptic prophages (monocins) in Listeria and sequence analysis of a holin/endolysin gene. Microbiology 141, 2577–2584 (1995).

20. Lee, G. et al. F-Type Bacteriocins of *Listeria monocytogenes*: a New Class of Phage Tail-Like Structures Reveals Broad Parallel Coevolution between Tailed Bacteriophages and High-Molecular-Weight Bacteriocins. J. Bacteriol. 198, 2784–2793 (2016).

21. Sigal, N. et al. Specialized *Listeria monocytogenes* produce tailocins to provide a population-level competitive growth advantage. Nat. Microbiol. 9, 2727–2737 (2024).

22. Gu, Z., Ge, X. & Wang, J. Structure of an F-type phage tail-like bacteriocin from *Listeria monocytogenes*. Nat. Commun. 16, 1695 (2025).

23. Gebhart, D. et al. Novel High-Molecular-Weight, R-Type Bacteriocins of *Clostridium difficile*. J. Bacteriol. 194, 6240–6247 (2012).

24. Cai, X. et al. Atomic structures of a bacteriocin targeting Gram-positive bacteria. Nat. Commun. 15, 7057 (2024).

25. Cheng, S. et al. Complete Genome Sequence of a New *Enterococcus faecalis* Bacteriophage, vB_EfaS_IME197. Genome Announc. 4, e00827–16 (2016).

26. Edgar, R. C. MUSCLE: multiple sequence alignment with high accuracy and high throughput. Nucleic Acids Res. 32, 1792–1797 (2004).

27. Afonina, I. et al. The composition and function of *Enterococcus faecalis* membrane vesicles. microLife 2, uqab002 (2021).

28. McBride, S. M., Fischetti, V. A., LeBlanc, D. J., Moellering, R. C. & Gilmore, M. S. Genetic Diversity among Enterococcus faecalis. PLoS ONE 2, e582 (2007).

29. Powell, H. R., Islam, S. A., David, A. & Sternberg, M. J. E. Phyre2.2: A Community Resource for Template-based Protein Structure Prediction. J. Mol. Biol. 437, 168960 (2025).

30. Dunne, M. et al. Reprogramming Bacteriophage Host Range through Structure-Guided Design of Chimeric Receptor Binding Proteins. Cell Rep. 29, 1336–1350.e4 (2019).

31. Abramson, J. et al. Accurate structure prediction of biomolecular interactions with AlphaFold 3. Nature 630, 493–500 (2024).

32. Spinelli, S., Veesler, D., Bebeacua, C. & Cambillau, C. Structures and host-adhesion mechanisms of lactococcal siphophages. Front. Microbiol. 5, (2014).

33. Ayala, R. et al. Nearly complete structure of bacteriophage DT57C reveals architecture of head-to-tail interface and lateral tail fibers. Nat. Commun. 14, 8205 (2023).

34. d’Acapito, A., Decombe, A., Arnaud, C.-A. & Breyton, C. Comparative anatomy of siphophage tails before and after interaction with their receptor. Curr. Opin. Struct. Biol. 92, 103045 (2025).

35. Xu, Y., Murray, B. E. & Weinstock, G. M. A Cluster of Genes Involved in Polysaccharide Biosynthesis from *Enterococcus faecalis* OG1RF. Infect. Immun. 66, 4313–4323 (1998).

36. Teng, F., Singh, K. V., Bourgogne, A., Zeng, J. & Murray, B. E. Further Characterization of the *epa* Gene Cluster and Epa Polysaccharides of *Enterococcus faecalis*. Infect. Immun. 77, 3759–3767 (2009).

37. Palmer, K. L. et al. Comparative Genomics of Enterococci: Variation in *Enterococcus faecalis*, Clade Structure in E. faecium, and Defining Characteristics of *E. gallinarum* and *E. casseliflavus*. mBio 3, e00318–11 (2012).

38. Salamzade, R. et al. zol and fai: large-scale targeted detection and evolutionary investigation of gene clusters. Nucleic Acids Res. 53, gkaf045 (2025).

39. Clewell, D. B. et al. Mapping of *Streptococcus faecalis* plasmids pAD1 and pAD2 and studies relating to transposition of Tn917. J. Bacteriol. 152, 1220–1230 (1982).

40. Sharpe, M. E. Serological Types of *Streptococcus faecali*s and its Varieties and their Cell Wall Type Antigen. J. Gen. Microbiol. 36, 151–160 (1964).

41. Thurlow, L. R., Thomas, V. C. & Hancock, L. E. Capsular Polysaccharide Production in *Enterococcus faecalis* and Contribution of CpsF to Capsule Serospecificity. J. Bacteriol. 191, 6203–6210 (2009).

42. Guerardel, Y. et al. Complete Structure of the Enterococcal Polysaccharide Antigen (EPA) of Vancomycin-Resistant *Enterococcus faecalis* V583 Reveals that EPA Decorations Are Teichoic Acids Covalently Linked to a Rhamnopolysaccharide Backbone. mBio 11, e00277–20 (2020).

43. Davis, J. L. et al. Dissecting the Enterococcal Polysaccharide Antigen (EPA) structure to explore innate immune evasion and phage specificity. Carbohydr. Polym. 347, 122686 (2025).

44. Gilmore, M. S. & Haas, W. The selective advantage of microbial fratricide. Proc. Natl. Acad. Sci. 102, 8401–8402 (2005).

45. Chatterjee, A. et al. Bacteriophage Resistance Alters Antibiotic-Mediated Intestinal Expansion of Enterococci. Infect. Immun. 87, e00085–19 (2019).

46. Lebreton, F. et al. Emergence of Epidemic Multidrug-Resistant *Enterococcus faecium* from Animal and Commensal Strains. mBio 4, e00534–13 (2013).

47. Schloissnig, S. et al. Genomic variation landscape of the human gut microbiome. Nature 493, 45–50 (2013).

48. Clewell, D. B. et al. Extrachromosomal and Mobile Elements in Enterococci: Transmission, Maintenance, and Epidemiology. in Enterococci: From Commensals to Leading Causes of Drug Resistant Infection (eds Gilmore, M. S., Clewell, D. B., Ike, Y. & Shankar, N.) (Massachusetts Eye and Ear Infirmary, Boston, 2014).

49. Ghequire, M. G. K. & De Mot, R. The Tailocin Tale: Peeling off Phage Tails. Trends Microbiol. 23, 587–590 (2015).

50. Krupovic, M. & Koonin, E. V. Multiple origins of viral capsid proteins from cellular ancestors. Proc. Natl. Acad. Sci. 114, (2017).

51. Sarris, P. F., Ladoukakis, E. D., Panopoulos, N. J. & Scoulica, E. V. A Phage Tail-Derived Element with Wide Distribution among Both Prokaryotic Domains: A Comparative Genomic and Phylogenetic Study. Genome Biol. Evol. 6, 1739–1747 (2014).

52. Punjani, A., Rubinstein, J. L., Fleet, D. J. & Brubaker, M. A. cryoSPARC: algorithms for rapid unsupervised cryo-EM structure determination. Nat. Methods 14, 290–296 (2017).

53. Posit team (2025). RStudio: Integrated Development Environment for R. Posit Software, PBC, Boston, MA.

54. Kutter, E. Phage Host Range and Efficiency of Plating. in Bacteriophages (eds Clokie, M. R. J. & Kropinski, A. M.) vol. 501 141–149 (Humana Press, Totowa, NJ, 2009).

55. Carlson, K. Working with Bacteriophages: Common Techniques and Methodological Approaches, Vol 1. CRC Press, Boca Raton, FL.

56. Broad Institute. Data from Bioproject PRJNA313452. NCBI Genbank.

57. Broad Institute. Data from Bioproject PRJNA663900. NCBI Genbank.

58. Mass Eye and Ear/Harvard Medical School. NCBI Genbank. Data from Bioproject PRJNA1371623.

59. Li, H. & Durbin, R. Fast and accurate short read alignment with Burrows–Wheeler transform. Bioinformatics 25, 1754–1760 (2009).

60. Walker, B. J. et al. Pilon: An Integrated Tool for Comprehensive Microbial Variant Detection and Genome Assembly Improvement. PLoS ONE 9, e112963 (2014).

61. Darling, A. E. et al. PhyloSift: phylogenetic analysis of genomes and metagenomes. PeerJ 2, e243 (2014).

62. Parks, D. H., Imelfort, M., Skennerton, C. T., Hugenholtz, P. & Tyson, G. W. CheckM: assessing the quality of microbial genomes recovered from isolates, single cells, and metagenomes. Genome Res. 25, 1043–1055 (2015).

63. Jain, C., Rodriguez-R, L. M., Phillippy, A. M., Konstantinidis, K. T. & Aluru, S. High throughput ANI analysis of 90K prokaryotic genomes reveals clear species boundaries. Nat. Commun. 9, 5114 (2018).

64. Van Dijk, L. R. et al. StrainGE: a toolkit to track and characterize low-abundance strains in complex microbial communities. Genome Biol. 23, 74 (2022).

65. Arndt, D. et al. PHASTER: a better, faster version of the PHAST phage search tool. Nucleic Acids Res. 44, W16–W21 (2016).

66. Seemann, T. Prokka: rapid prokaryotic genome annotation. Bioinformatics 30, 2068–2069 (2014).

67. Mistry, J. et al. Pfam: The protein families database in 2021. Nucleic Acids Res. 49, D412–D419 (2021).

68. Kanehisa, M. KEGG: Kyoto Encyclopedia of Genes and Genomes. Nucleic Acids Res. 28, 27–30 (2000).

69. Karp, P. D. et al. The BioCyc collection of microbial genomes and metabolic pathways. Brief. Bioinform. 20, 1085–1093 (2019).

70. Price, M. N. & Arkin, A. P. PaperBLAST: Text Mining Papers for Information about Homologs. mSystems 2, e00039–17 (2017).

71. Georgescu, C. H. et al. SynerClust: a highly scalable, synteny-aware orthologue clustering tool. *Microb*. Genomics 4, (2018).

72. Schrödinger, L. & DeLano, W. PyMOL. (2020).

73. Pedregosa, F. et al. Scikit-learn: Machine learning in Python. Journal of Machine Learning Research 12, 2825–2830 (2011).

74. Emms, D. M. & Kelly, S. OrthoFinder: phylogenetic orthology inference for comparative genomics. Genome Biol. 20, 238 (2019).

75. Katoh, K., Kuma, K., Toh, H. & Miyata, T. MAFFT version 5: improvement in accuracy of multiple sequence alignment. 8.

76. Minh, B. Q. et al. IQ-TREE 2: New Models and Efficient Methods for Phylogenetic Inference in the Genomic Era. Mol. Biol. Evol. 37, 1530–1534 (2020).

77. Gilmore, S. M., Earl, A., Prichula, J., Manson, A, Richards, G., Karumidze, N. & Lebreton, F. Compositions and methods for control of specific bacterial populations. US Patent WO2025054631, filed September 10, 2023. https://patentscope.wipo.int/search/en/WO2025054631.

78. Lossouarn, J. et al. Enterococcus faecalis Countermeasures Defeat a Virulent Picovirinae Bacteriophage. Viruses 11, 48 (2019).

