## Supplementary Information for "Efagins – engineerable agents that evolved independently to target the enterococcal cell wall"

**This file includes:**

- Supplementary Figures – S1-S6
- Supplementary Word Tables: Table S26 and S27

### Supplementary Figures

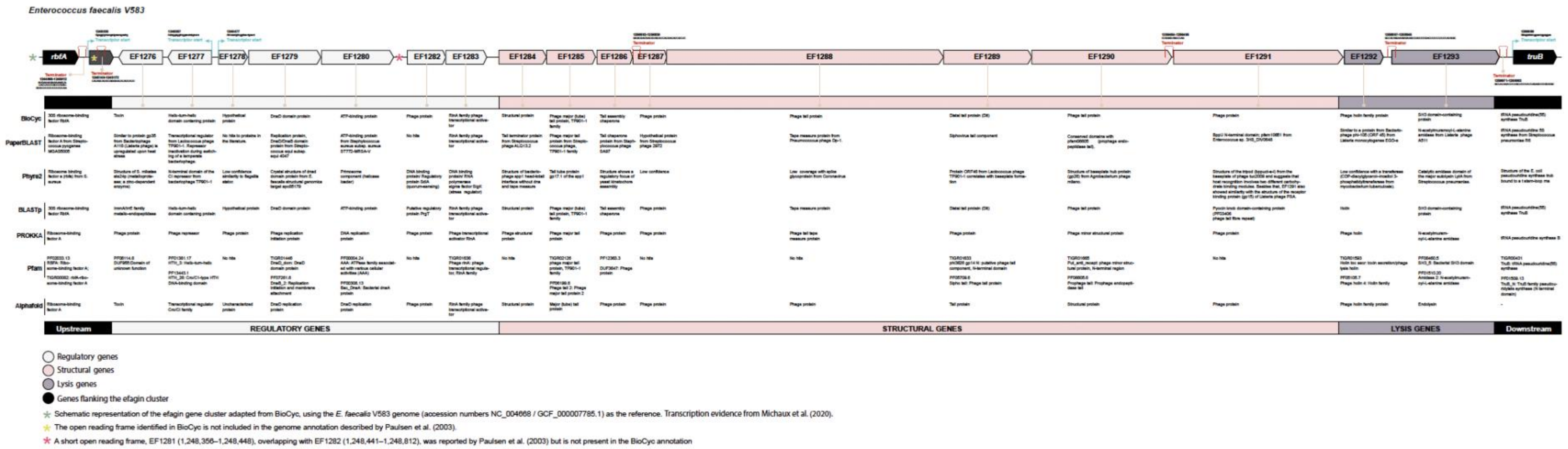


**Supplementary Fig. 1. V583 efagin gene function prediction**. Biocyc,[1] PaperBlast,[2] Phyre2.2,[3] Blastp,[4] Prokka,[5] Pfam,[6] and Alphafold[7] were used to infer the function.

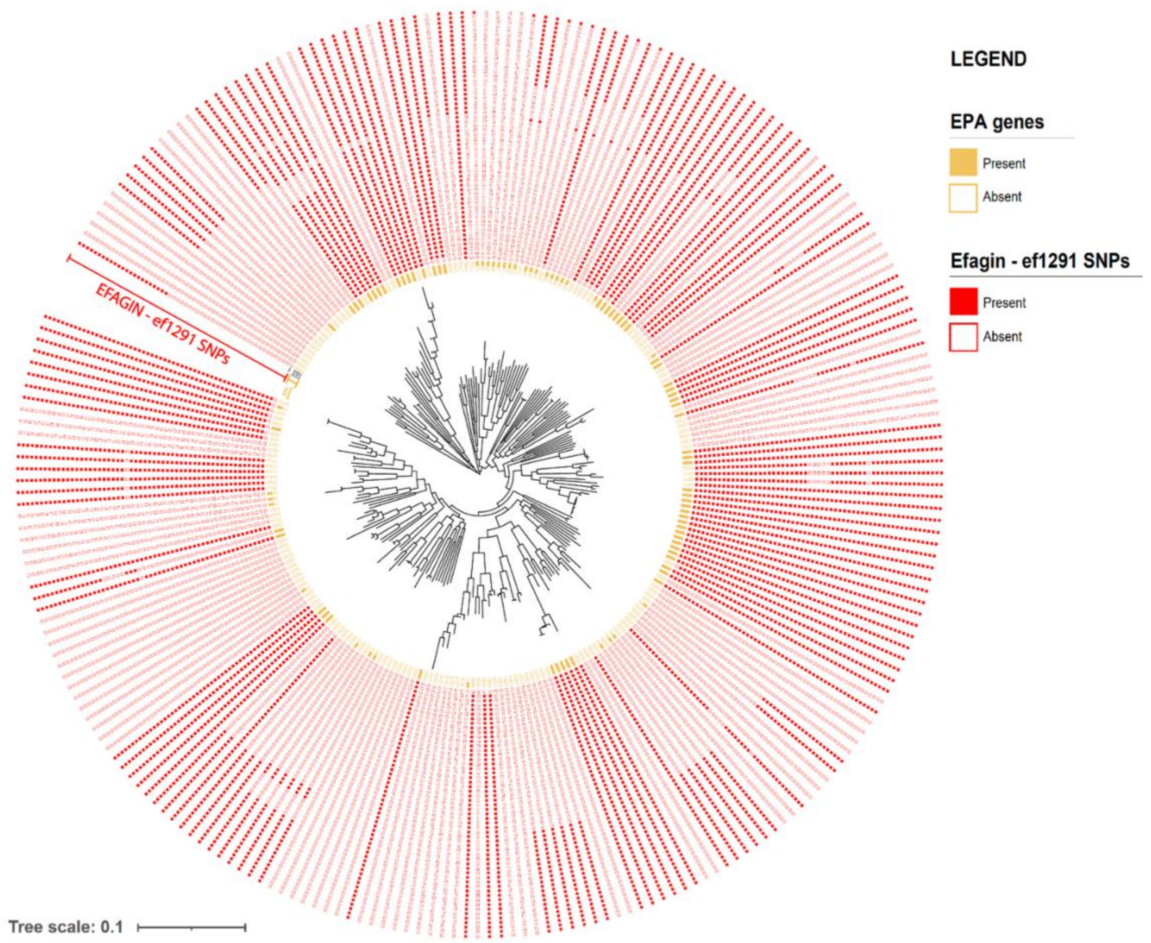


**Supplementary Fig. 2. Variation of *epa* gene presence/absence and efagin-associated SNPs across *E. faecalis* representative genomes.** This plot shows the distribution of *epa* gene presence (yellow) and efagin *ef*1291 SNPs (red) across *E. faecalis* strains, aligned to a whole-genome phylogeny (center). Concordance between these *ef*1291 SNPs and the presence of these specific *epa* genes was observed in 81% of strains, far exceeding the ~50% expected by chance, suggesting coordinated evolutionary dynamics. The enrichment of these reciprocal patterns supports a functional or selective relationship between efagin and the *epa* locus.

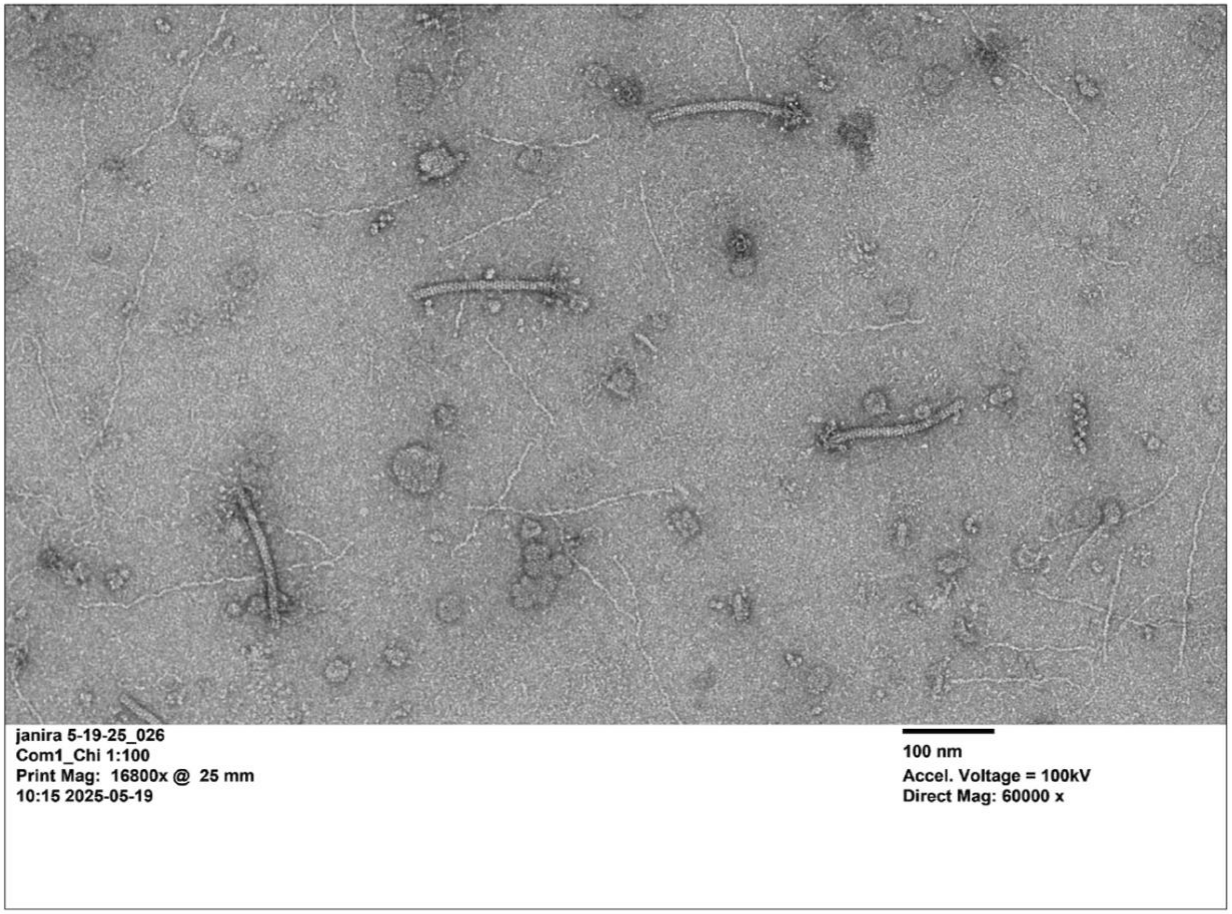


**Supplementary Fig. 3. Visualization of efagin from OG1RF*χefgD* chimera strain.** Transmission electron microscopy (TEM) shows images of efagin particles produced by the *E. faecalis* OG1RF*χefgD* chimera strain expressing type D efagin. Morphological features were visualized and quantified using TEM and analyzed with Fiji/ImageJ software.

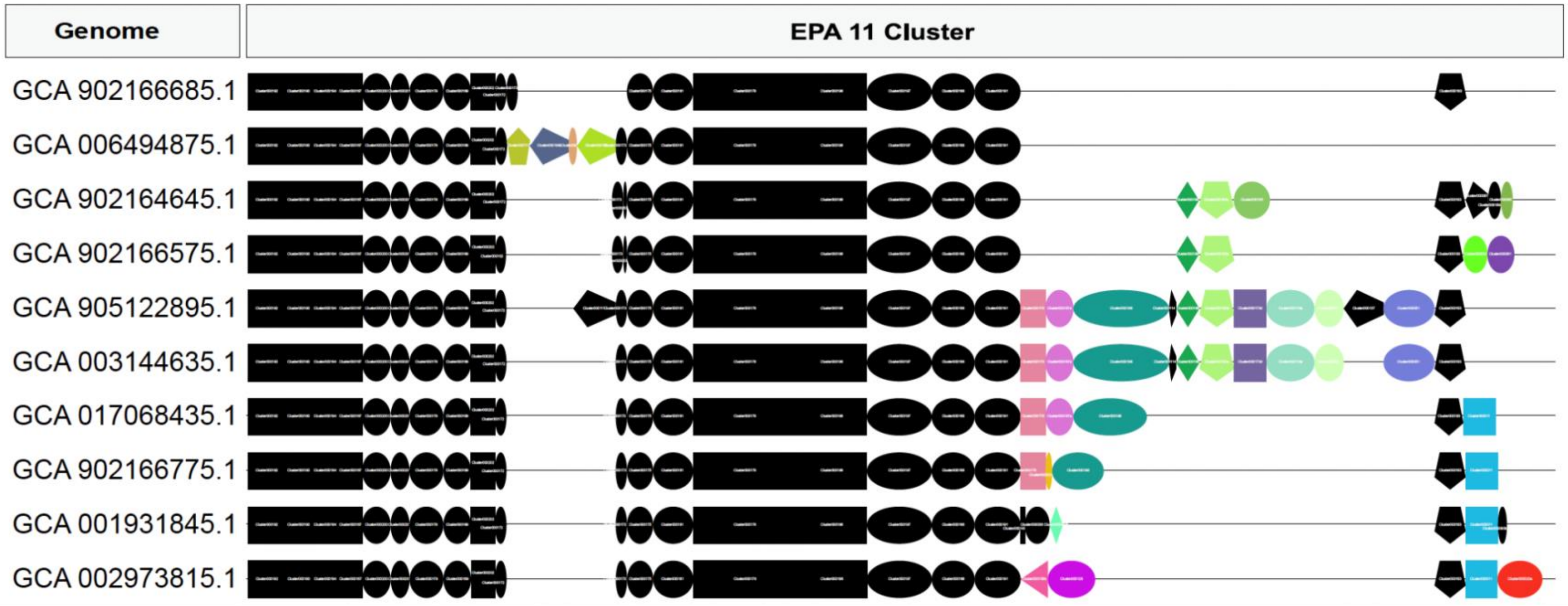


**Supplementary Fig. 4. Gene structure of representatives of *Epa* type 11 in 10 *E. faecalis* genomes**. Each row represents a distinct *E. faecalis* genome, labeled by its NCBI accession number. Conserved core *epa* genes are shown in black, while variable genes (e.g., 000192, 000190, etc.) are represented in distinct colors, highlighting differences in the epa operon across strains, within *Epa* type 11.

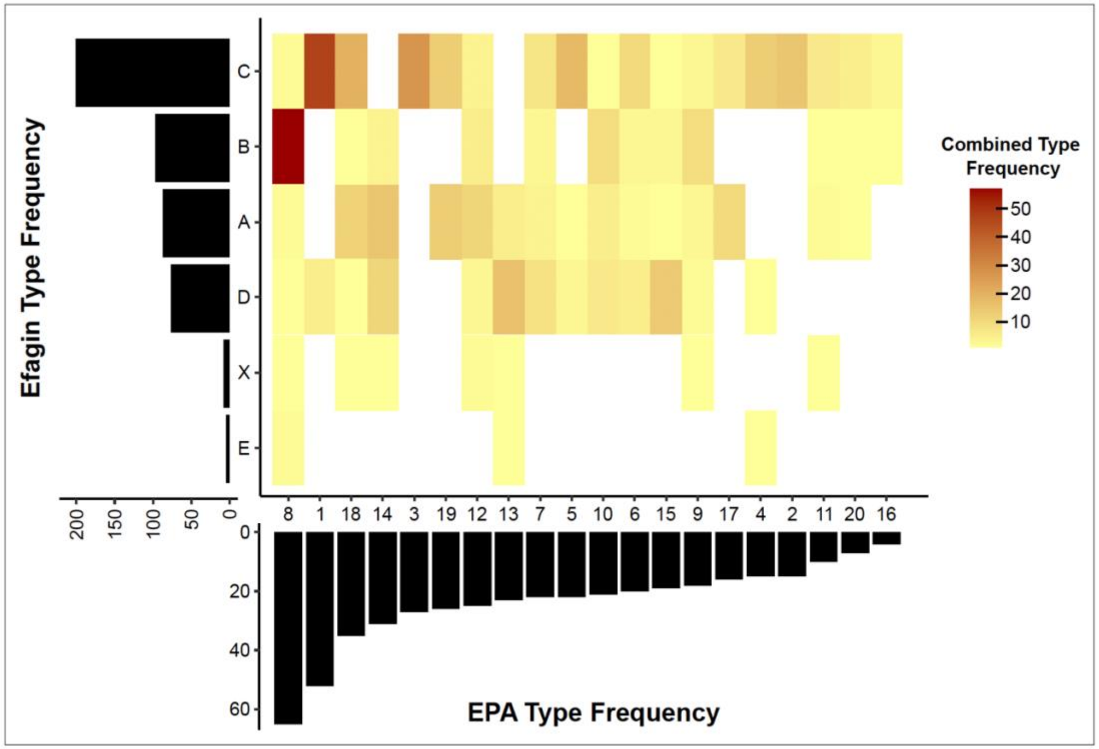


**Supplementary Fig. 5. Frequency of combinations of efagin and EPA types present in 493 *E. faecalis* representative genomes.** Rows represent different efagin types; columns represent different *epa* types. The central heat map indicates the frequency of each combination of *epa* and efagin types; marginal bar plots show the frequency of each *epa* or efagin type individually across the 493 genomes. A white square indicates that there are no genomes with that combination of efagin and *epa* types.

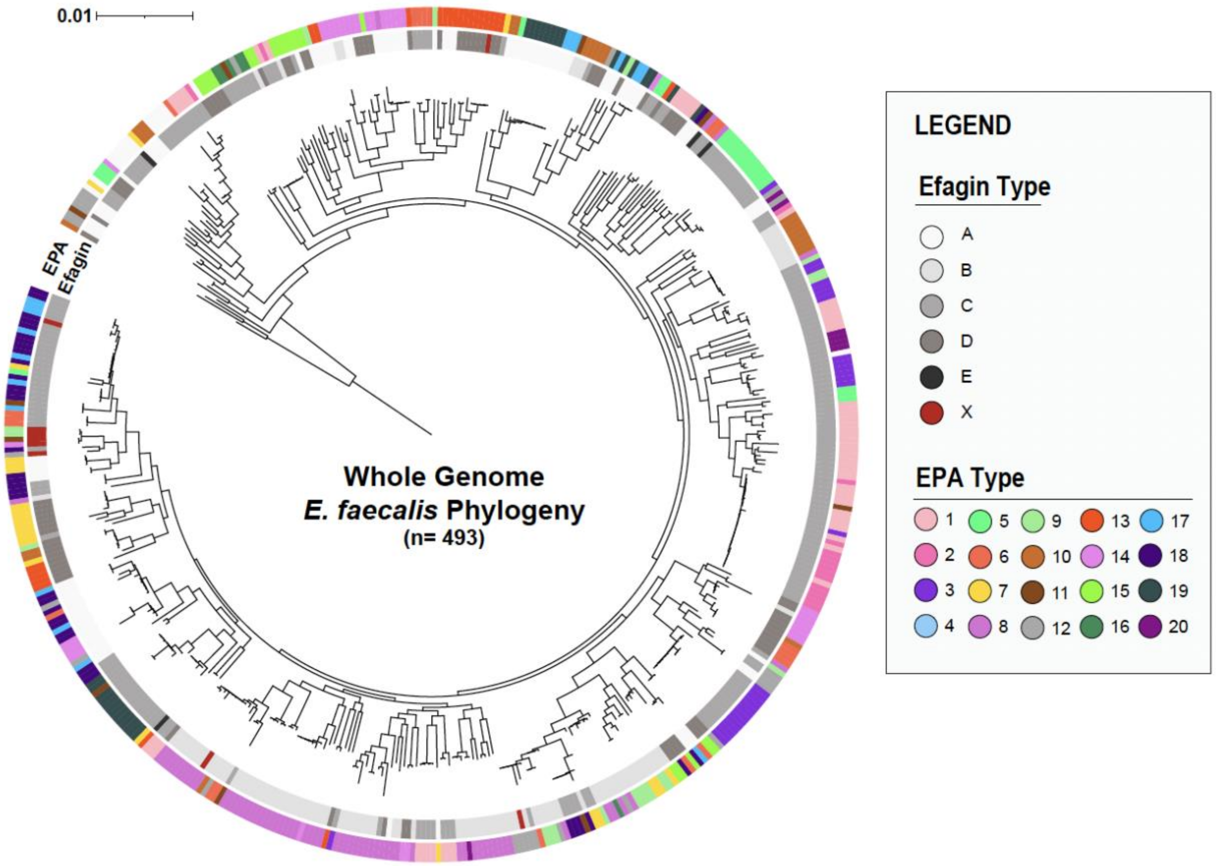


**Supplementary Fig. S6. Whole-genome phylogeny of *E. faecalis* strains, overlaid with the diversity of *epa* types (outer ring; 1-20) and efagin types (inner ring; A-E, and X).** The lack of concordance between the genomic phylogeny and the distribution of efagin or epa types suggests that efagin and *epa* genetic diversity were likely acquired by horizontal gene transfer and localized recombination. Scale bar indicates genetic distance.

**Supplementary Word Tables**

**Table S26. List of strains used in this study**

| Strains | Species | Description | Source |
|---|---|---|---|
| OG1RF | *E. faecalis* | Oral isolate selected for spontaneous resistances to rifampicin and fusidic acid | Dunny et al.[8] |
| OG1RF*Δefg* | *E. faecalis* | OG1RF with 14.360 bp of 14.6kb of efagin locus deleted | **This work** |
| *OG1RF*χefgA* | *E. faecalis* | OG1RF chimera with type A efagin *ef1291* domain from donor *E. faecalis* T9 | **This work** |
| *OG1RF*χefgB* | *E. faecalis* | OG1RF chimera with type B efagin *ef1291* domain from donor from donor *E. faecalis* T6 | **This work** |
| *OG1RF*χefgD* | *E. faecalis* | OG1RF chimera with type D efagin *ef1291* domain from donor *E. faecalis* Com 1 | **This work** |
| *OG1RF*χefgE* | *E. faecalis* | OG1RF chimera with type E efagin *ef1291* domain from donor *E. faecalis* T19 strain. | **This work** |
| OG1RF-R1 | *E. faecalis* | OG1RF spontaneous mutant resistant to efagin type D. Deletion (1 bp) in *epa* decoration gene (OG1RF11711, Glycerol-phosphate transferase) | **This work** |
| OG1RF-R2 | *E. faecalis* | OG1RF spontaneous mutant resistant to efagin type D. Mutation (C->A; Gly-87-Val; NSY) in *epa* decoration gene (OG1RF11715 (*epa*OX); Glycosyl transferase, group 2 family protein) | **This wor**k |
| V583 | *E. faecalis* | Clinical strain | Sahm et al.[9] Paulsen et al.[10] |
| V583-R1 | *E. faecalis* | V583 spontaneous mutant resistant to efagin types A, B, D, and E. Insertion (657 bp) in *epa* decoration gene (EF2170 (*epa*X); glycosyl transferase, group 2 family protein) | **This work** |
| V583-R2 | *E. faecalis* | V583 spontaneous mutant resistant to efagin types A, B, D, and E. Insertion (321 bp) in *epa* decoration gene (EF2170 (*epa*X); glycosyl transferase, group 2 family protein) | **This work** |
| FA2-2 | *E. faecalis* | Spontaneous rifampicin and fusidic acid resistant derivative of clinical isolate | Clewell et al.[11] |
| FA2-2-R1 | *E. faecalis* | FA2-2 spontaneous mutant resistant to efagin types C and E. Deletion (2 bp) in UCI02299 (O-antigen ligase family protein) | **This work** |
| FA2-2-R2 | *E. faecalis* | FA2-2 spontaneous mutant resistant to efagin types C and E. Deletion (2 bp) in UCI02299 (O-antigen ligase family protein) | **This work** |
| FA2-2-R3 | *E. faecalis* | FA2-2 spontaneous mutant resistant to efagin types C and E. Mutation (C->A; Gly-377-Val; NSY) in UCI02299 (O-antigen ligase family protein) | **This work** |

| | | | |
|---|---|---|---|
| FA2-2-R4 | *E. faecalis* | FA2-2 spontaneous mutant resistant to efagin types C and E. Deletion (1 bp) in UCI02299 (O-antigen ligase family protein) | **This work** |
| X98 | *E. faecalis* | Historic pediatric clinical isolate | Sharpe et al.[12] |
| X98-R1 | *E. faecalis* | X98 spontaneous mutant resistant to efagin type C. Deletion (1 bp) in *epa* decoration gene EFOG00558 (oligosaccharide repeat unit polymerase) | **This work** |
| X98-R2 | *E. faecalis* | X98 spontaneous mutant resistant to efagin type C. Mutation (C->A; stop codon) in *epa* decoration gene EFOG00558 (oligosaccharide repeat unit polymerase) | **This work** |
| 51RS1 | *E. faecalis* | X98 spontaneous phage resistant mutant resistant to efagin type C. Deletion (24 bp) in *epa* decoration gene EFOG00558 (oligosaccharide repeat unit polymerase) | Chatterjee et al.[13] |
| 51RS2 | *E. faecalis* | X98 spontaneous phage resistant mutant resistant to efagin types A, C, and E. Mutation (C->T; Arg-295-Trp; NSY) in *epa* decoration gene EFOG00552 (glycosyl transferase family 2 protein) | Chatterjee et al.[13] |
| EC1000 | *Escherichia coli* | Laboratory strain. Plasmid free | - |
| EC1000 (pLT06) | *Escherichia coli* | Contains pLT06 plasmid | Thurlow et al.[14] |
| JM110 | Escherichia coli | Indicator strain used in the M13 plaque assay | - |
| Mp18 | *Escherichia coli* | M13 bacteriophage | - |

*Chimera is abbreviated using the Greek letter "χ"(chi), and efagin is abbreviated as "*efg*"

**Table S27. List of primers used in this study**

| Gene | Oligo name | Sequence 5' to 3' | Size (bp) | T(C) | Reference |
|---|---|---|---|---|---|
| *ef*1291 **(Efagin Chimeras)** | 817_F_BamHI | GGTGGT**GGATCC**AATCAACGTGTGAAAGCCGC **BamHI** | 1325 | 60 | **This work** |
| | 2141_R_pstI | GGTGGT**CTGCAG**GGCATAGGAATAAAGGTCGC **pstI** | | | |
| *ef*1291 **(Efagin A)** | 1542F_A | CCACATGCCTAAAGTATTTAAC | 109 | 60 | **This work** |
| | 1650R_A | ATACGGAAAAACGTAAGGTGC | | | |
| *ef*1291 **(Efagin B)** | 1541F_B | CCCACATGCCTAAAATATT | 126 | 53 | **This work** |
| | 1669R_B | GTATCCTAATTGTAATGCAC | | | |
| *ef*1291 **(Efagin C)** | 1575F_C | ATATACATTATCGACGAAGATTA | 106 | 58 | **This work** |
| | 1680R_C | TCCCATCTCATATAAACCTGC | | | |
| *ef*1291 **(Efagin D)** | 1575F_D | TTATAGTATAAATTCTGTTCTC | 106 | 56 | **This work** |
| | 1680R_D | ATCTATTTCGTACAAGGCTGT | | | |
| *ef*1291 **(Efagin E)** | 1343F_E | CCGAAGCTCCTTTATTCAAAA | 386 | 57 | **This work** |
| | 1728R_E | TATGACCATTCCAAGTCCAT | | | |
| **Efagin deletion** | KO-F1_XbaI | GTGGT**TCTAGA**GTCCAAGGCATCACTATTAC **XbaI** | 878 | 58 | **This work** |
| | KO-R1_Acc65I | CGC**GGTACC**AGTTGAAAGCCCGCAATGTG **Acc65I** | | | |
| **Efagin deletion** | KO-F2_KpnI | GCG**GGTACC**ATACATGGGGATCATTTACTAG **KpnI** | 860 | 59 | **This work** |
| | KO-R2-SbfI | GTGGT**CCTGCAGG**GATTGTTTCAGCAGCCATTTG | | | |

| | | | | | |
|---|---|---|---|---|---|
| | | **SbfI** | | | |
| **Efagin deletion** | Internal_F | ATTCCTCGGCTAAAATGTTC | 619 | 59 | **This work** |
| | Internal_R | CTGGCATCGGTAAATCTTC | | | |
| **pLT06** | OriF | CAATAATCGCATCCGATTGCA | - | 56 | Thurlow et al.[14] |
| | KS05seqR | CCTATTATACCATATTTTGGAC | | | |

[14]Primers used for the allelic replacement.

.

### References


1. Karp, P. D. *et al.* The BioCyc collection of microbial genomes and metabolic pathways. *Brief. Bioinform.* **20**, 1085–1093 (2019).

2. Price, M. N. & Arkin, A. P. PaperBLAST: Text Mining Papers for Information about Homologs. *mSystems* **2**, e00039-17 (2017).

3. Powell, H. R., Islam, S. A., David, A. & Sternberg, M. J. E. Phyre2.2: A Community Resource for Template-based Protein Structure Prediction. *J. Mol. Biol.* **437**, 168960 (2025).

4. National Center for Biotechnology Information (NCBI).

5. Seemann, T. Prokka: rapid prokaryotic genome annotation. *Bioinformatics* **30**, 2068–2069 (2014).

6. Mistry, J. *et al.* Pfam: The protein families database in 2021. *Nucleic Acids Res.* **49**, D412–D419 (2021).

7. Abramson, J. *et al.* Accurate structure prediction of biomolecular interactions with AlphaFold 3. *Nature* **630**, 493–500 (2024).

8. Dunny, G. M., Brown, B. L. & Clewell, D. B. Induced cell aggregation and mating in Streptococcus faecalis: evidence for a bacterial sex pheromone. *Proc. Natl. Acad. Sci.* **75**, 3479–3483 (1978).

9. Sahm, D. F. *et al.* In vitro susceptibility studies of vancomycin-resistant Enterococcus faecalis. *Antimicrob. Agents Chemother.* **33**, 1588–1591 (1989).

10. Paulsen, I. T. Role of Mobile DNA in the Evolution of Vancomycin-Resistant Enterococcus faecalis. *Science* **299**, 2071–2074 (2003).

11. Clewell, D. B. *et al.* Mapping of Streptococcus faecalis plasmids pAD1 and pAD2 and studies relating to transposition of Tn917. *J. Bacteriol.* **152**, 1220–1230 (1982).

12. Sharpe, M. E. Serological Types of Streptococcus faecalis and its Varieties and their Cell Wall Type Antigen. *J. Gen. Microbiol.* **36**, 151–160 (1964).

13. Chatterjee, A. *et al.* Bacteriophage Resistance Alters Antibiotic-Mediated Intestinal Expansion of Enterococci. *Infect. Immun.* **87**, e00085-19 (2019).

14. Thurlow, L. R., Thomas, V. C. & Hancock, L. E. Capsular Polysaccharide Production in *Enterococcus faecalis* and Contribution of CpsF to Capsule Serospecificity. *J. Bacteriol.* **191**, 6203–6210 (2009).
