## Supplementary material for "Efagins – engineerable agents that evolved independently to target the enterococcal cell wall": Graphical Abstract

### EFAGINS

*E. faecalis* phage-related inhibitors

Conserved *E. faecalis* genomes

Target include VRE

Five natural variants identified

Programmable to target EPA variations

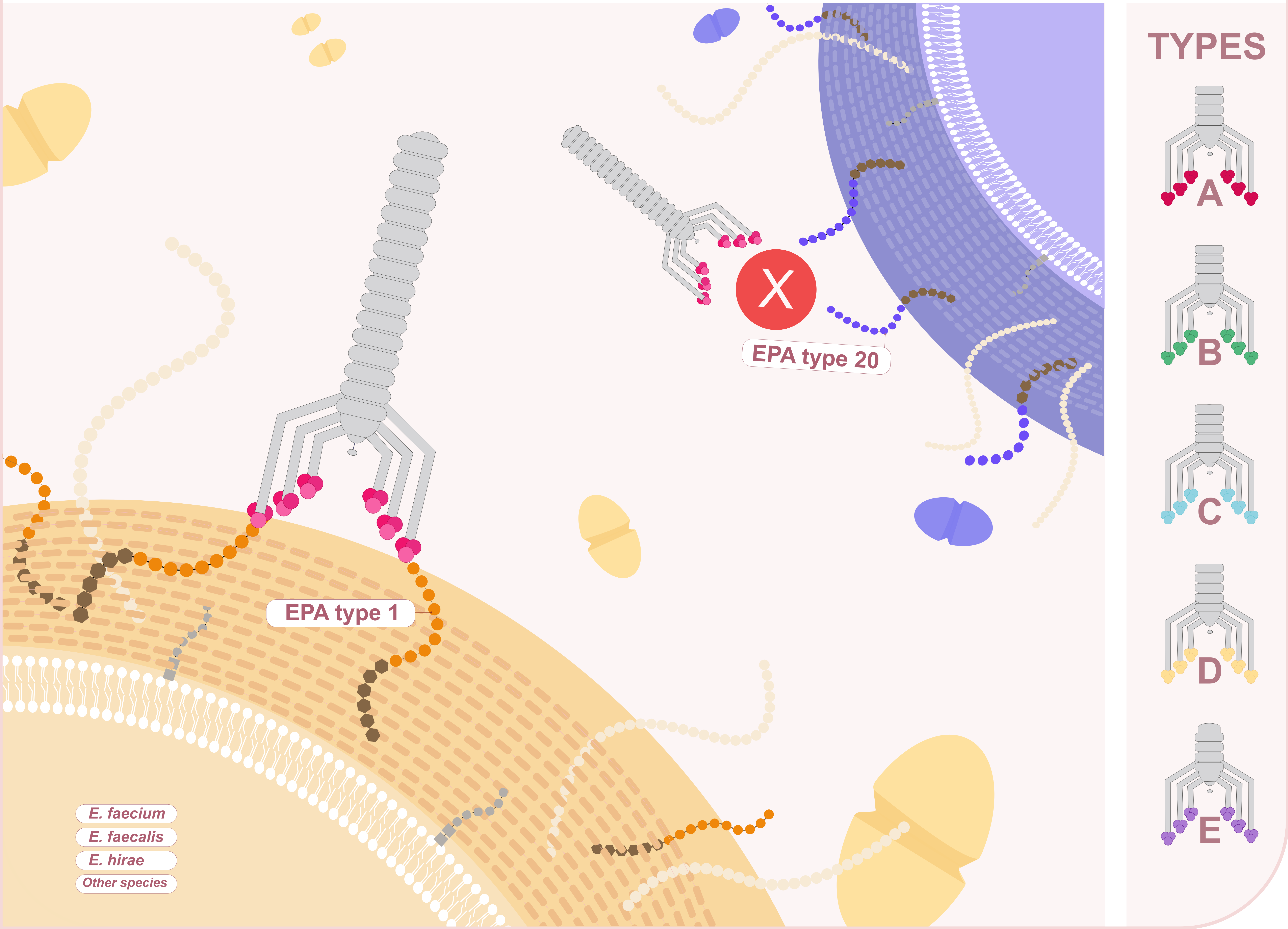
