## Supplementary material for "Efagins – engineerable agents that evolved independently to target the enterococcal cell wall": Highlights

- Identified Efagins – *E. faecalis* phage-related inhibitors – conserved and encoded within all *E. faecalis* genomes
- Targeting specificity is driven by polymorphisms in tail fiber–like proteins and cell wall polysaccharide diversity
- Efagins inhibit many species of enterococci, including multidrug-resistant enterococci, vancomycin-resistant *E. faecium* and *E. faecalis* strains
- Efagins are programmable entities that can selectively target drug-resistant hospital clones while sparing commensal microbes
