## Extended Data Figures 1-10 for "Efagins – engineerable agents that evolved independently to target the enterococcal cell wall"

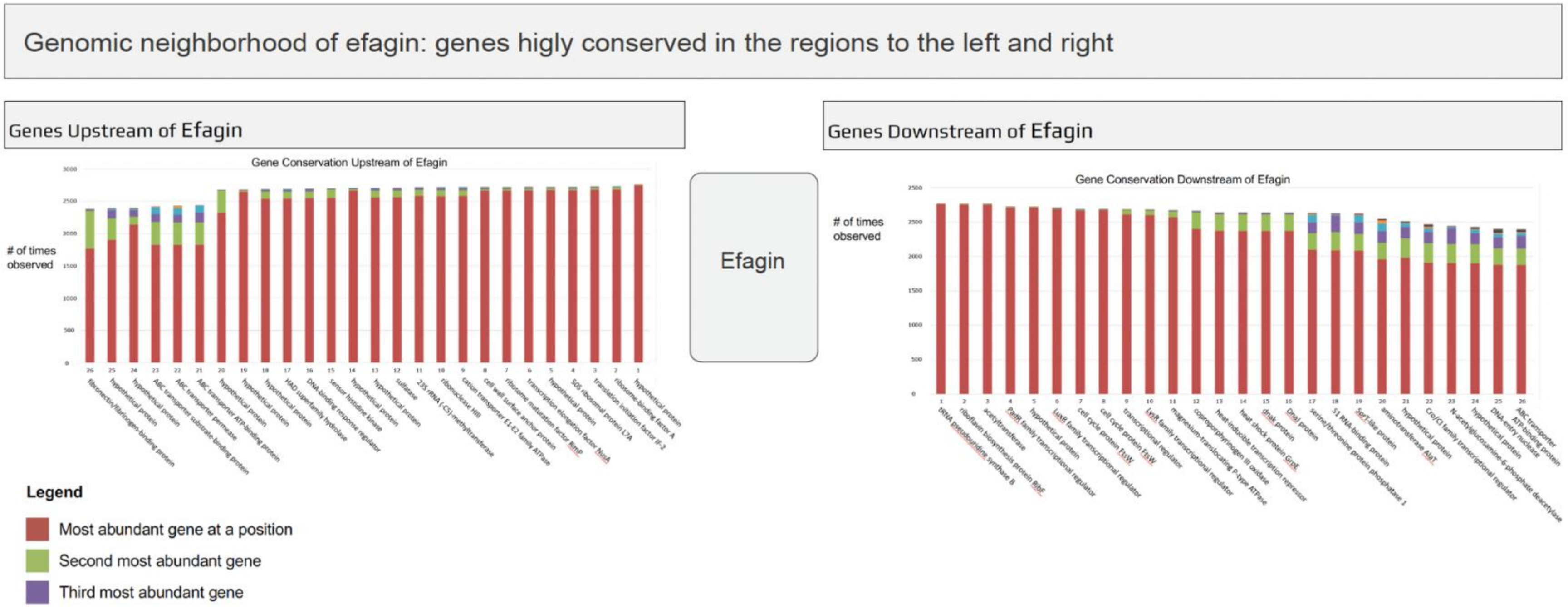


**Extended Data Fig. 1. Genomic neighborhood of efagin shows conserved flanking genes**. Bar plots display the frequency of genes found upstream (left) and downstream (right) of efagin across *E. faecalis* genomes. The most common gene at each position is shown in red, the second in green, and the third in purple.

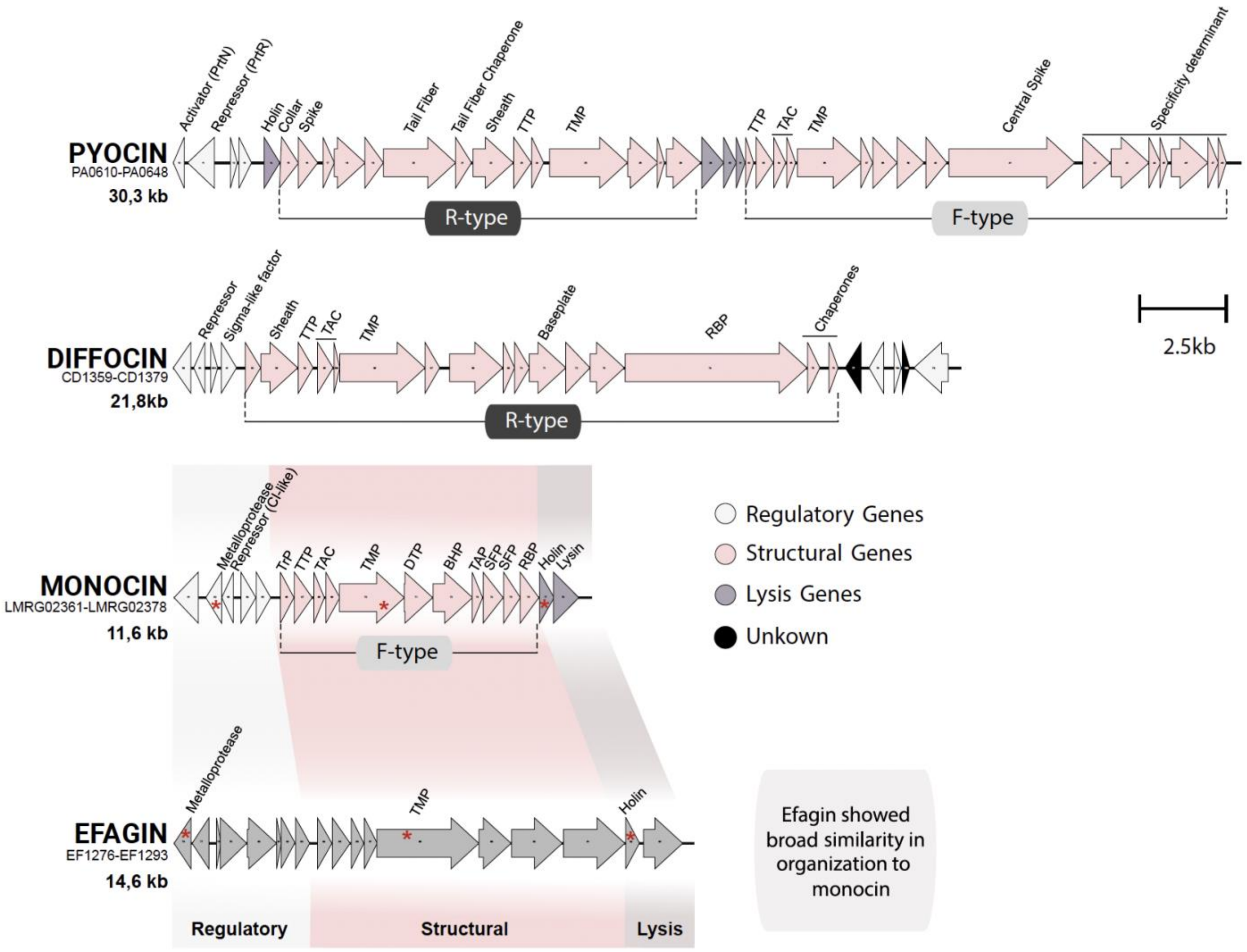
PYOCIN
PA0610-PA0648
30,3 kb
Activator (PrtN)
Repressor (PrtR)
Holin
Collar
Spike
Tail Fiber
Tail Fiber Chaperone
Sheath
TTP
TMP
TTP
TAC
TMP
Central Spike
Specificity determinant
R-type
F-type
DIFFOCIN
CD1359-CD1379
21,8kb
Repressor
Sigma-like factor
Sheath
TTP
TAC
TMP
Baseplate
RBP
Chaperones
R-type
2.5kb
MONOCIN
LMRG02361-LMRG02378
11,6 kb
Metalloprotease
Repressor (CI-like)
TrP
TTP
TAC
TMP
DTP
BHP
TAP
SFP
SFP
RBP
Holin
Lysin
F-type
Regulatory Genes
Structural Genes
Lysis Genes
Unkown
EFAGIN
EF1276-EF1293
14,6 kb
Metalloprotease
TMP
Holin
Regulatory
Structural
Lysis
Efagin showed broad similarity in organization to monocin

**Extended Data Fig. 2. Comparison of gene clusters encoding phage tail-like elements.** Pyocin (30.3 kb), diffocin (21.8 kb), monocin (11.6 kb), and efagin (14.6 kb) from *Pseudomonas aeruginosa*, *Clostridium difficile*, *Listeria monocytogenes*, and *Enterococcus faecalis*, respectively. While these systems show broad similarity in gene organization, they share little to no sequence identity, suggesting independent evolutionary origins. Genomic comparison between the *L. monocytogenes* monocin locus and the *E. faecali*s efagin locus reveals similar gene organization, particularly in structural modules such as the tube, tape measure, baseplate components, and receptor-binding protein (RBP). Although efagin shares organizational similarity with monocin, it has minimal sequence identity with monocin across three genes (red asterisk). Both systems encode non-contractile phage tail-like structures and appear to have arisen independently. Arrows represent open reading frames (ORFs), color-coded by predicted function: regulatory (lightly grey), structural (pink), and lysis-associated genes (purple). Particles are classified as R-type (contractile) or F-type (non-contractile). Reference sequences used include: i) the R- and F-type pyocin clusters from *Pseudomonas aeruginosa* PAO1, corresponding to the genomic region spanning genes PA0610–PA0648 (Genbank: AE004091.2); ii) the R-type diffocin from *Clostridium difficile* 630 (GenBank: NZ_CP010905.2; genes CD1359–CD1379); iii) the F-type monocin from *Listeria monocytogenes* 10403S (GenBank: NC_017544.1; genes LMRG02361–LMRG02378); and iv) efagin from *E. faecalis* V583 (GenBank: NC_004668.1; genes ef1276–ef1293) were included as reference sequences. The number inside each arrow denotes the number of genes represented.

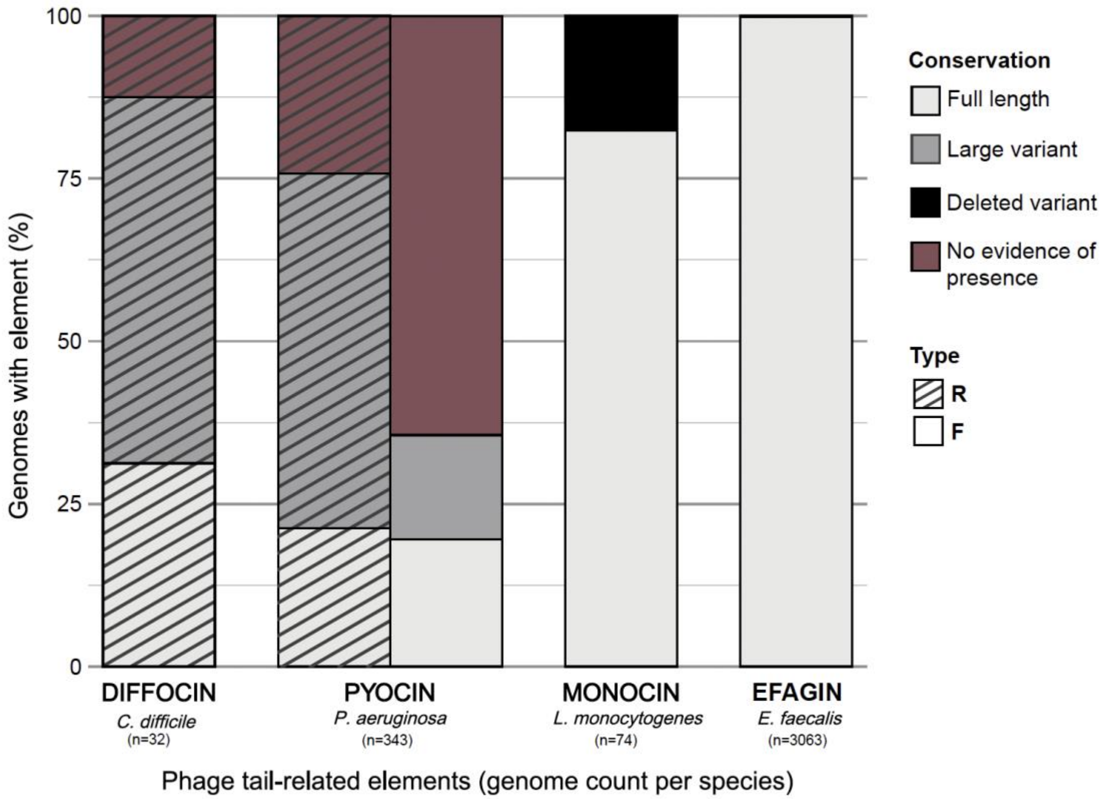


**Extended Data Fig. 3. Distribution of contractile (R-type) and non-contractile (F-type) phage tail-like elements among genomes from Gram-positive and Gram-negative bacterial species.** In contrast to efagin, which is highly conserved and intact in all strains for which high-quality genome sequences are available, the other tailocins appear variable and often occur in alternate hybrid forms.

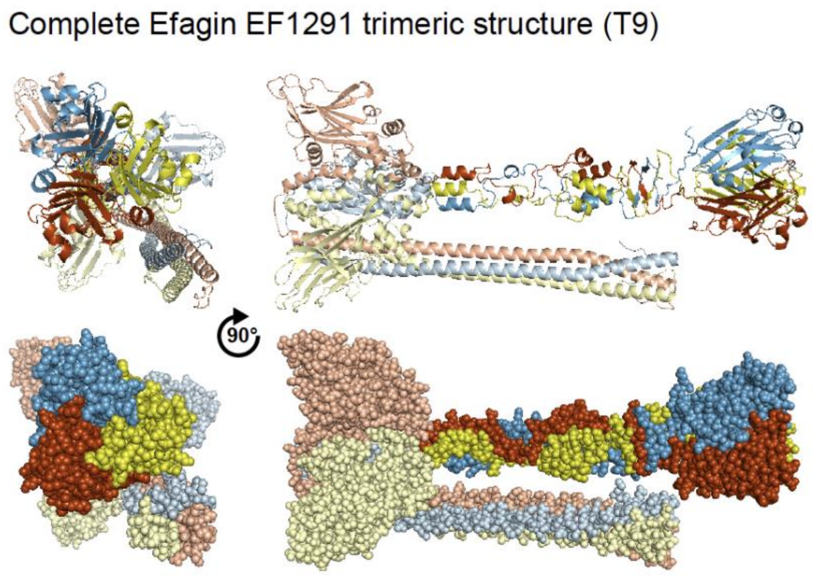


**Extended Data Fig. 4. Predicted model for EF1291 as ribbon and space-filling structures using Alphafold 3.**[31] The three colors (blue, red, and yellow) represent individual monomers within the trimer. Regions highlighted with darker colors in the C-terminal end correspond to the EF1291 structural regions related to the receptor-binding domain of Listeria phage PSA protein gp15 by Phyre2.2.[29]

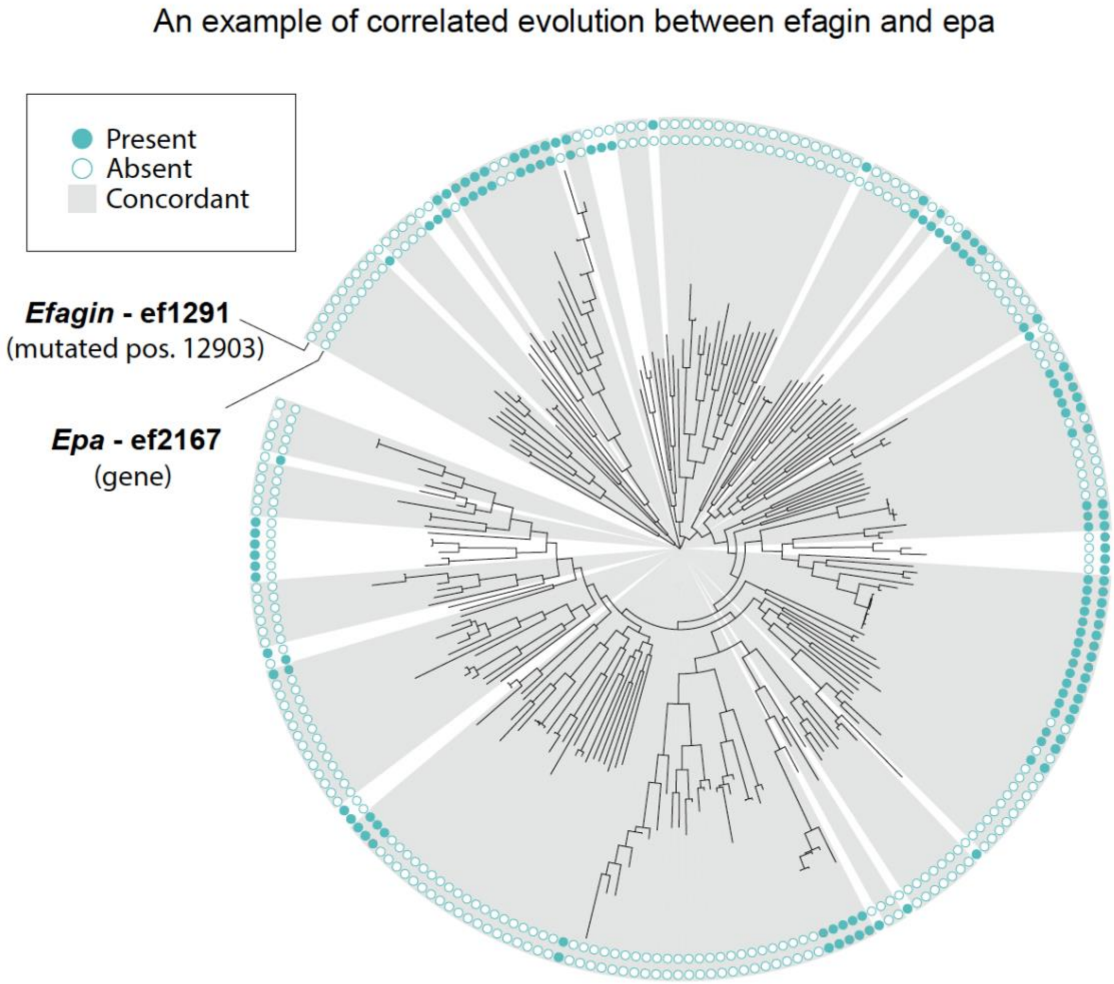


**Extended Data Fig. 5. Coordinated variation of *epa* gene presence/absence and efagin-associated SNPs across *E. faecalis* representative genomes.** Example of a correlation between a sequence variant in efagin *ef*1291 and the presence or absence of a specific gene in the *epa,* across the diversity of *E. faecalis* strains. This plot shows variation in position 12903 of *ef*1291 and the presence or absence of the *epa* gene *ef*2167. Gray shading indicates presence or absence concordance between mutated position 12903 and *ef*2167. Similar correlations were observed between efagin positions and *ef*2166, *ef*2167, *ef*2168, and *ef*2169, four adjacent genes within the *epa*.

**a**

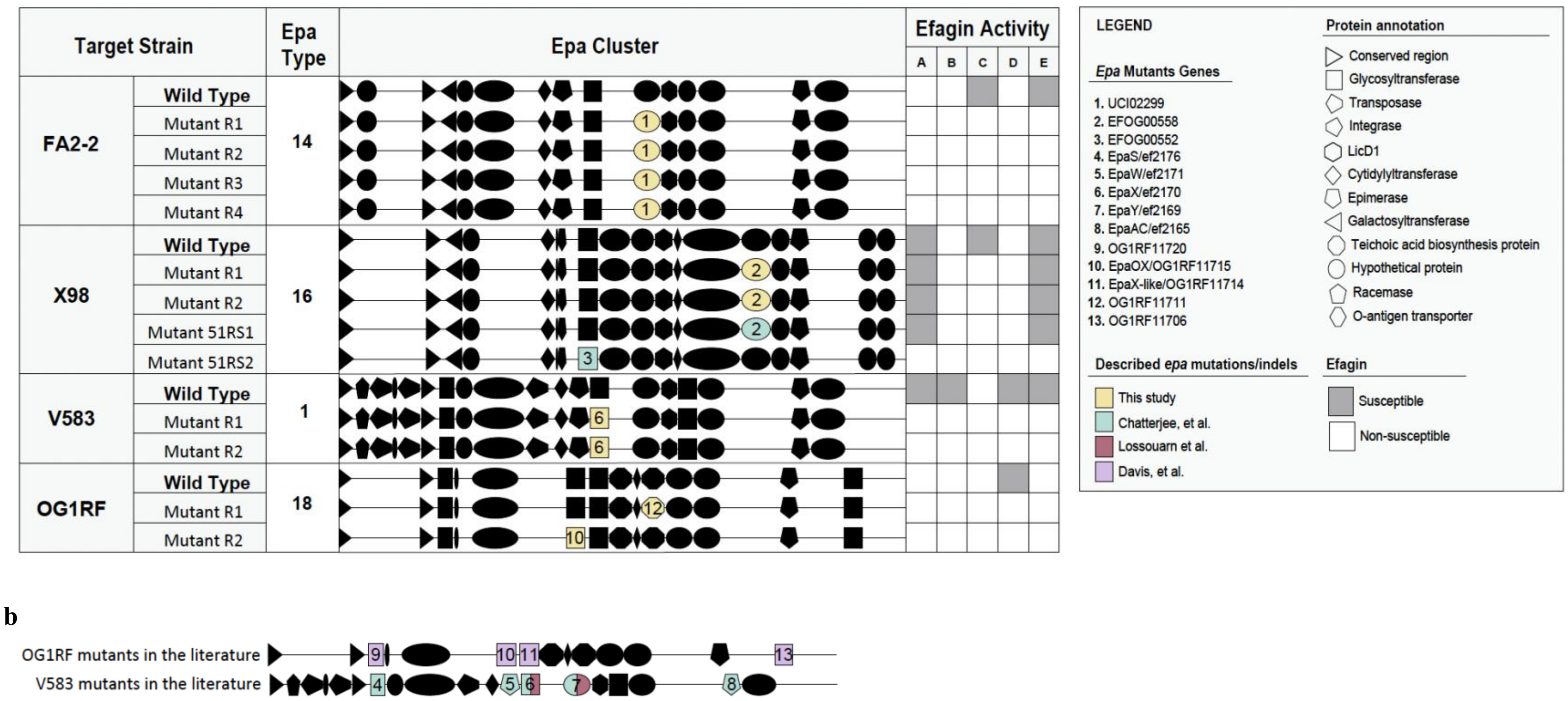


**Extended Data Fig. 6. Efagin resistant mutants have defects in *epa* variable region genes. a,** Effect of *epa* mutations on efagin susceptibility in four *E. faecalis* strains (FA2-2, X98, V583, and OG1RF). **b,** Other studies[43,45,78] describing *epa* mutations or indels correlated with phage resistance.

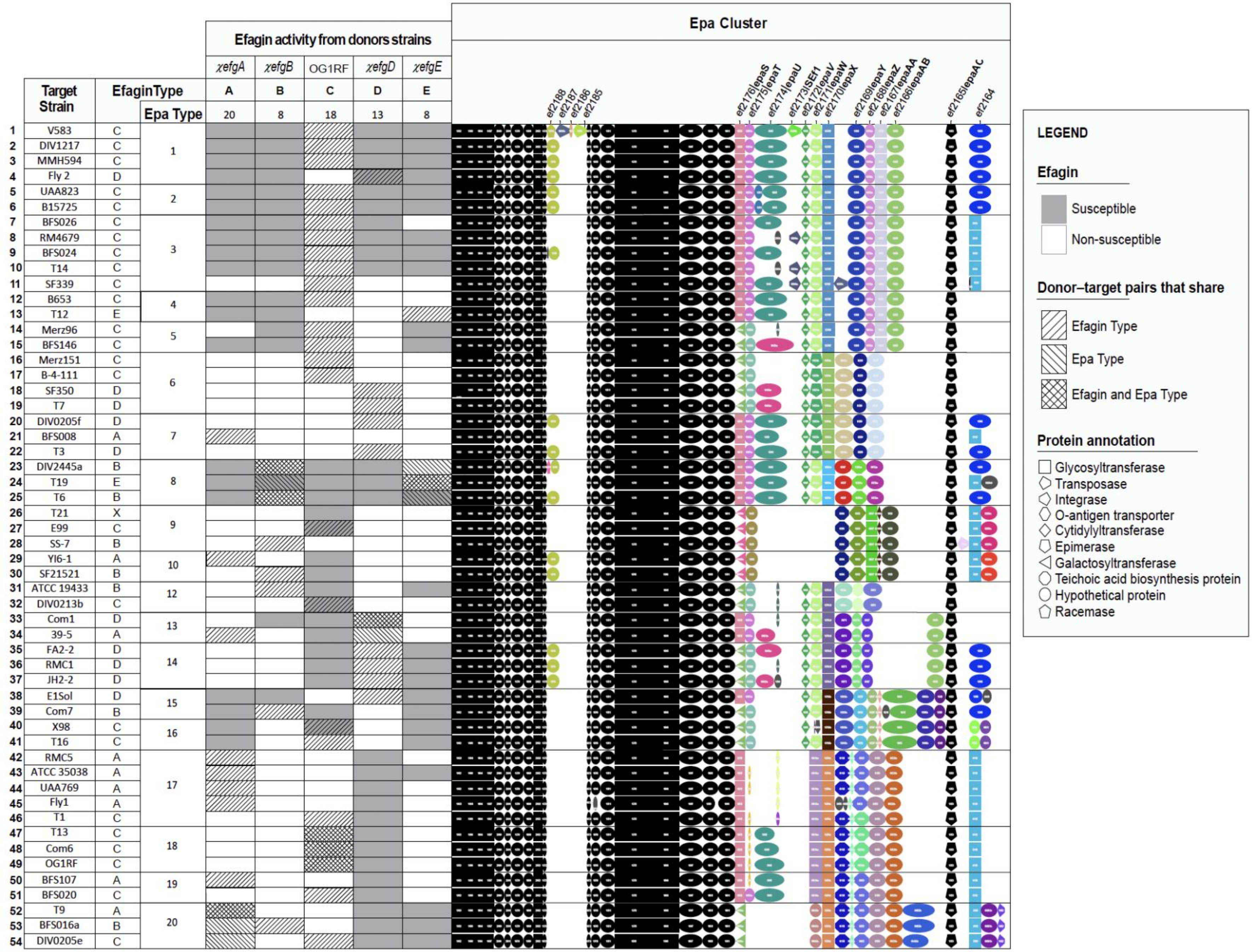
Efagin activity from donors strains
χefgA
χefgB
OG1RF
χefgD
χefgE
A B C D E
20 8 18 13 8
Epa Cluster
ef2188
ef2187
ef2186
ef2185
ef2176|epaS
ef2175|epaT
ef2174|epaU
ef2173|ISEF1
ef2172|epaV
ef2171|epaW
ef2170|epaX
ef2169|epaY
ef2168|epaZ
ef2167|epaAA
ef2166|epaAB
ef2165|epaAC
ef2164
Target Strain
EfaginType
Epa Type
1 V583 C 1
2 DIV1217 C
3 MMH594 C
4 Fly 2 D
5 UAA823 C 2
6 B15725 C
7 BFS026 C 3
8 RM4679 C
9 BFS024 C
10 T14 C
11 SF339 C
12 B653 C 4
13 T12 E
14 Merz96 C 5
15 BFS146 C
16 Merz151 C 6
17 B-4-111 C
18 SF350 D
19 T7 D
20 DIV0205f D 7
21 BFS008 A
22 T3 D
23 DIV2445a B 8
24 T19 E
25 T6 B
26 T21 X 9
27 E99 C
28 SS-7 B
29 YI6-1 A 10
30 SF21521 B
31 ATCC 19433 B 12
32 DIV0213b C
33 Com1 D 13
34 39-5 A
35 FA2-2 D 14
36 RMC1 D
37 JH2-2 D
38 E1Sol D 15
39 Com7 B
40 X98 C 16
41 T16 C
42 RMC5 A 17
43 ATCC 35038 A
44 UAA769 A
45 Fly1 A
46 T1 C
47 T13 C 18
48 Com6 C
49 OG1RF C
50 BFS107 A 19
51 BFS020 C
52 T9 A 20
53 BFS016a B
54 DIV0205e C
LEGEND
Efagin
Susceptible
Non-susceptible
Donor–target pairs that share
Efagin Type
Epa Type
Efagin and Epa Type
Protein annotation
Glycosyltransferase
Transposase
Integrase
O-antigen transporter
Cytidylyltransferase
Epimerase
Galactosyltransferase
Teichoic acid biosynthesis protein
Hypothetical protein
Racemase

**Extended Data Fig. 7. Relationship between efagin activity and *epa* operon architecture across 54 *E. faecalis* strains.** Schematic figure showing the left-hand heat maps display efagin activity assays from five different donor strain: OG1RF (type C efagin) and OG1RF chimeras: *χefgA* (type A efagin), *χefgB* (type B efagin), *χefgD* (type D efagin), and *χefgE* (type E efagin), annotated with its *epa* operon type and assigned efagin type (A-E) against a diverse collection of *E. faecalis* strains (n=54). Grey indicates susceptible and white non-susceptibility. Grey hashing highlights donor–target pairs that share either the same efagin (hashes pointing up to the right) or/ and *epa* type (hashes pointing up to the left), indicating situations where we would not expect inhibition, and most often do not observe inhibition (**Table S19**). Operon cluster IDs are listed for each target strain. The figure highlights how variations in *epa* gene content, and efagin type are associated with differential killing profiles. Chimeric constructs are denoted using the Greek letter "*χ*" (chi), and efagin is abbreviated as "*efg*."

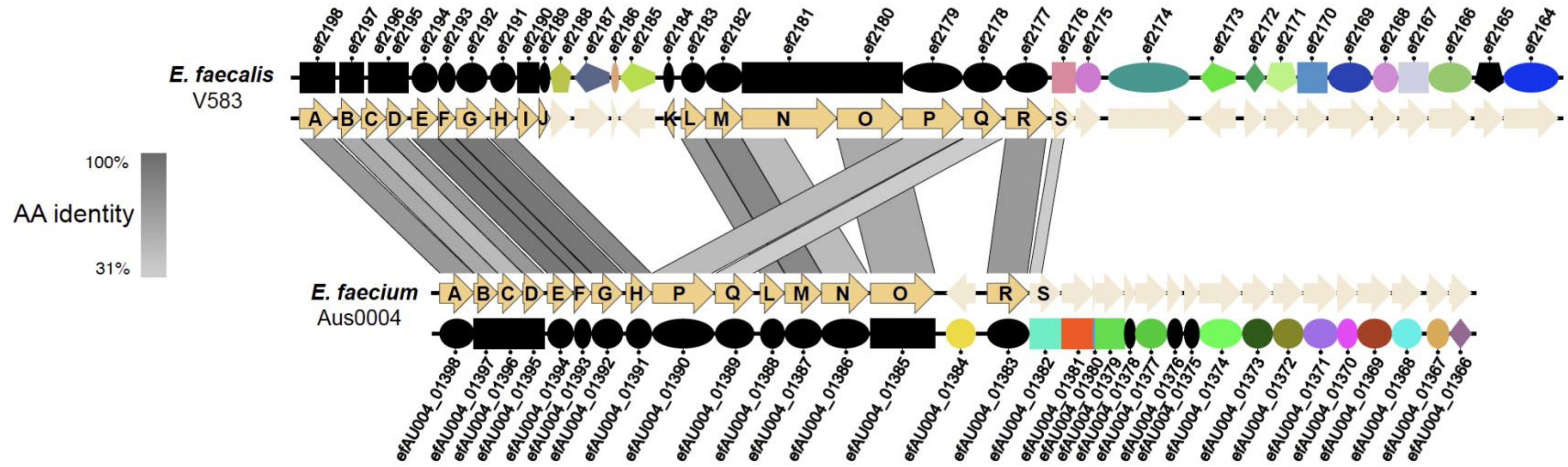


**Extended Data Fig. 8. Comparison of the organization of the *epa* locus between *E. faecalis* V583 and *E. faecium* Aus0004.** The *epa* rhamnose backbone is conserved in both species (dark yellow arrows), but differs in gene composition and arrangement, whereas the variable region (light yellow arrows) is completely different in the two species. Core *epa* genes (*epa*A–*epa*R) show moderate to high sequence identity (31.2–92.1%), whereas the variable decoration genes (light yellow) lack detectable homology, highlighting species-specific diversification of surface polysaccharide structures. In *E. faecium*, the locus follows the order *epa*ABCDEFGHPQLMNOR. The *E. faecium epa*N gene resembles *E. faecalis epa*N at the N-terminus, sharing a predicted S-adenosylmethionine binding site, but differs in its C-terminal sequence. We identified orthologs of *epa*P and *epa*Q in both species, but located at distinct genomic sites. In *E. faecalis*, the central region of the cluster contains *epa*I, *epa*J, and *epa*K, whereas in *E. faecium* these genes are absent and replaced by *epa*P and *epa*Q*,* suggesting the rhamnose polysaccharides of the two species have different structure.

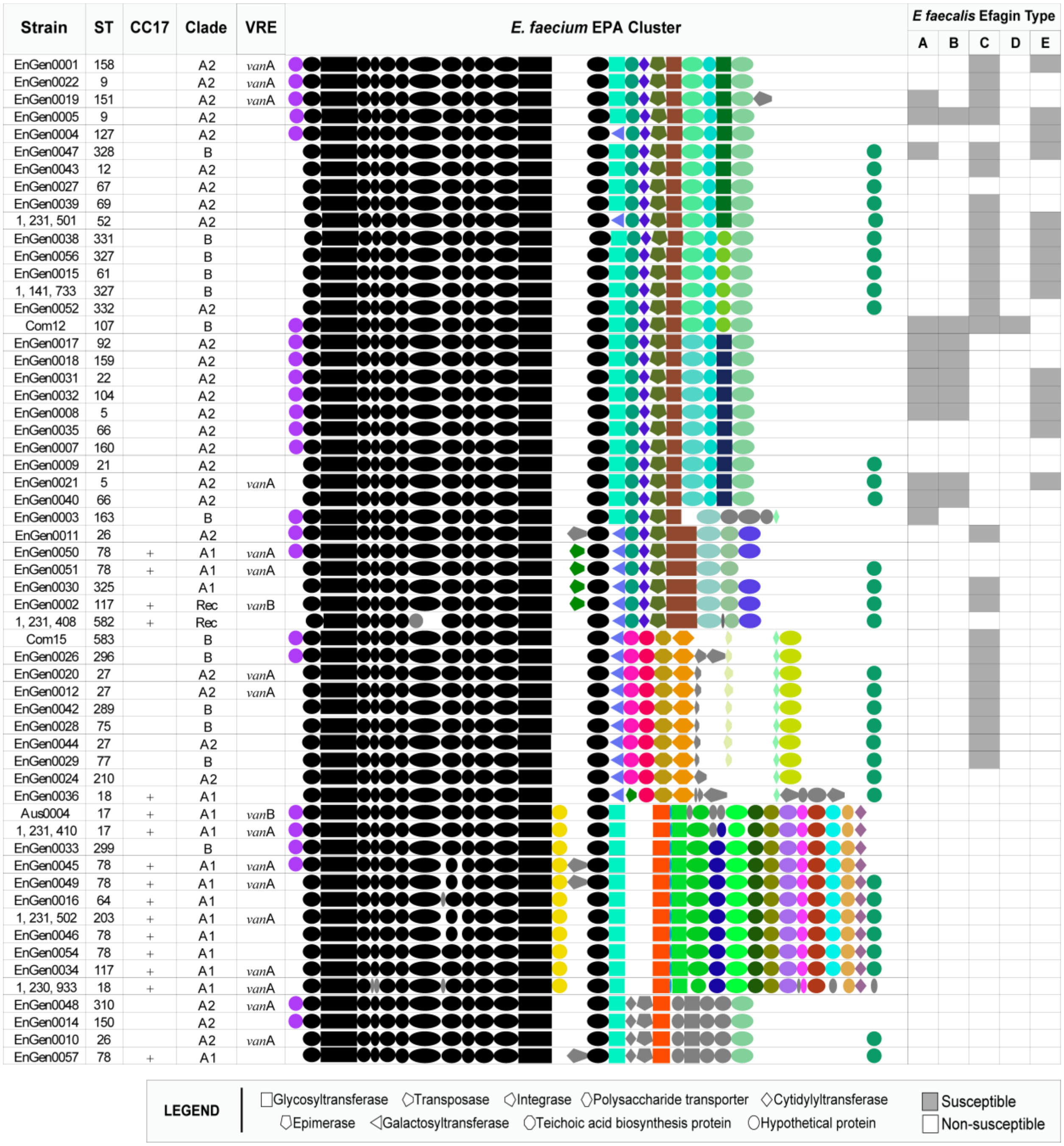


**Extended Data Fig. 9. Structural diversity of *epa* operons across *E. faecium* strains and correlation with efagin susceptibility**. Strain identifier, sequence type (ST), clonal complex (CC), *E. faecium* clade assignments (A1, A2, or B) and vancomycin resistance genotype in relation to the organization of *epa* operons. Rec indicates strains that are hybrids between clades. Conserved core genes are shown in black, while variable genes are colored according to orthogroup membership, with shapes corresponding to predicted function. The rightmost panel shows the efagin susceptibility profile. In the Epa cluster, black represents core genes found in all the genomes, and grey indicates rare genes found in less than 5%.

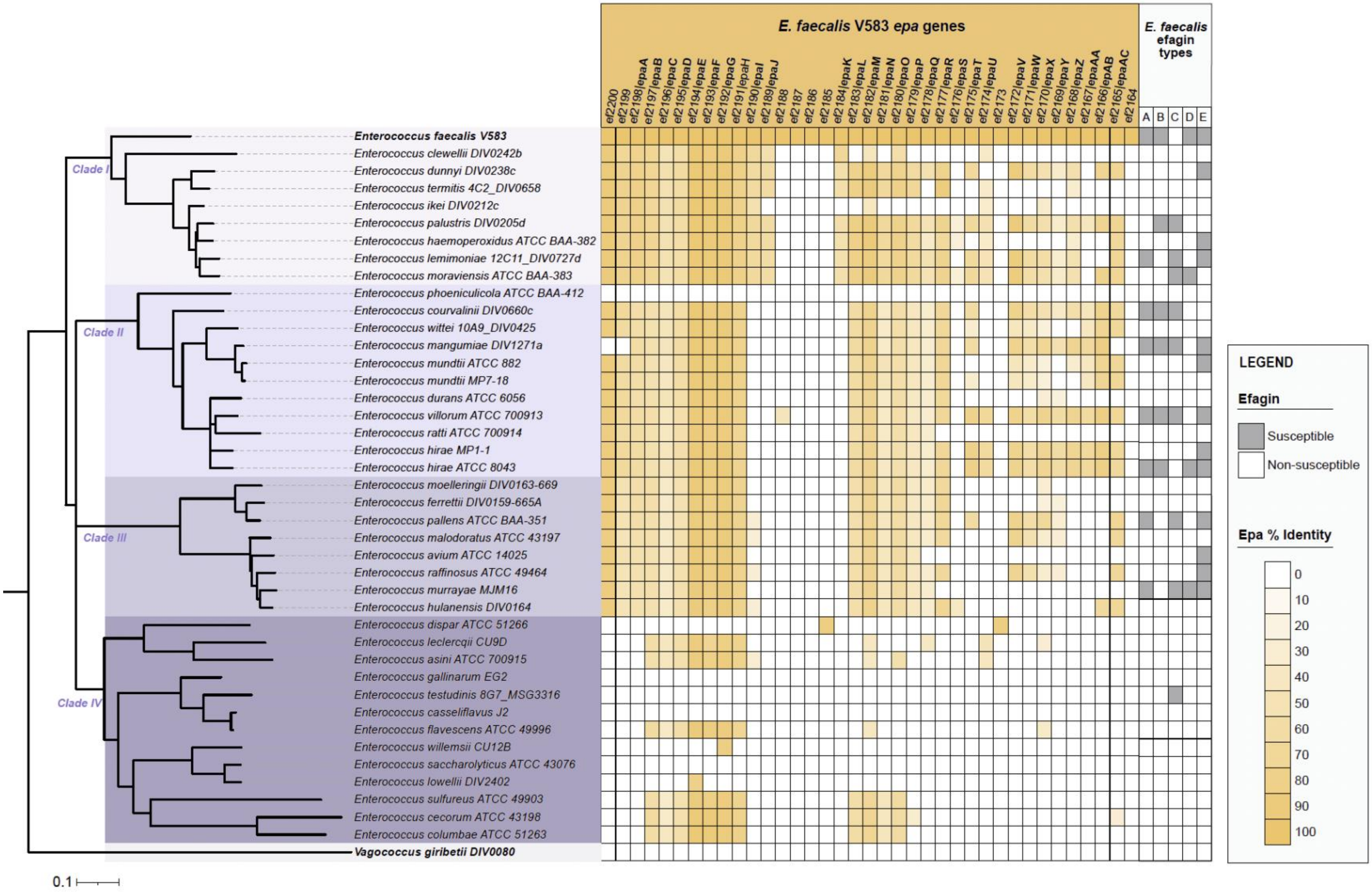
E. faecalis V583 epa genes
E. faecalis efagin types
ef2200
ef2199
ef2198|epaA
ef2197|epaB
ef2196|epaC
ef2195|epaD
ef2194|epaE
ef2193|epaF
ef2192|epaG
ef2191|epaH
ef2190|epaI
ef2189|epaJ
ef2188
ef2187
ef2186
ef2185
ef2184|epaK
ef2183|epaL
ef2182|epaM
ef2181|epaN
ef2180|epaO
ef2179|epaP
ef2178|epaQ
ef2177|epaR
ef2176|epaS
ef2175|epaT
ef2174|epaU
ef2173
ef2172|epaV
ef2171|epaW
ef2170|epaX
ef2169|epaY
ef2168|epaZ
ef2167|epaAA
ef2166|epaAB
ef2165|epaAC
ef2164
A
B
C
D
E
Clade I
Clade II
Clade III
Clade IV
Enterococcus faecalis V583
Enterococcus clewellii DIV0242b
Enterococcus dunnyi DIV0238c
Enterococcus termitis 4C2_DIV0658
Enterococcus ikei DIV0212c
Enterococcus palustris DIV0205d
Enterococcus haemoperoxidus ATCC BAA-382
Enterococcus lemimoniae 12C11_DIV0727d
Enterococcus moraviensis ATCC BAA-383
Enterococcus phoeniculicola ATCC BAA-412
Enterococcus courvalinii DIV0660c
Enterococcus wittei 10A9_DIV0425
Enterococcus mangumiae DIV1271a
Enterococcus mundtii ATCC 882
Enterococcus mundtii MP7-18
Enterococcus durans ATCC 6056
Enterococcus villorum ATCC 700913
Enterococcus ratti ATCC 700914
Enterococcus hirae MP1-1
Enterococcus hirae ATCC 8043
Enterococcus moelleringii DIV0163-669
Enterococcus ferrettii DIV0159-665A
Enterococcus pallens ATCC BAA-351
Enterococcus malodoratus ATCC 43197
Enterococcus avium ATCC 14025
Enterococcus raffinosus ATCC 49464
Enterococcus murrayae MJM16
Enterococcus hulanensis DIV0164
Enterococcus dispar ATCC 51266
Enterococcus leclercqii CU9D
Enterococcus asini ATCC 700915
Enterococcus gallinarum EG2
Enterococcus testudinis 8G7_MSG3316
Enterococcus casseliflavus J2
Enterococcus flavescens ATCC 49996
Enterococcus willemsii CU12B
Enterococcus saccharolyticus ATCC 43076
Enterococcus lowellii DIV2402
Enterococcus sulfureus ATCC 49903
Enterococcus cecorum ATCC 43198
Enterococcus columbae ATCC 51263
Vagococcus giribetii DIV0080
0.1
LEGEND
Efagin
Susceptible
Non-susceptible
Epa % Identity
0
10
20
30
40
50
60
70
80
90
100

**Extended Data Fig. 10. Conservation of *epa* operon genes across *Enterococcus* species and correlation with efagin susceptibility**. Strains of *Enterococcus* clades are colored in graded shades of purple by membership in clades I – IV.[1] The extent of amino acid sequence identity with inferred *Epa* gene products, when compared to the prototypes of *E. faecalis* strain V583, is indicated by the gold heatmap. Efagin susceptibility is indicated with grey shading.
